# When word order matters: human brains represent sentence meaning differently from large language models

**DOI:** 10.1101/2025.07.19.665701

**Authors:** James Fodor, Carsten Murawski, Shinsuke Suzuki

## Abstract

Large language models based on the transformer architecture are now capable of producing human-like language. But do they encode and process linguistic meaning in a human-like way? Here, we address this question by analysing 7T fMRI data from 30 participants reading 108 sentences each. These sentences are carefully designed to disentangle sentence structure from word meaning, thereby testing whether transformer representations of sentence meaning resemble those formed by the brain. We found that while transformer models match brain representations better than models that completely ignore word order, all transformer models performed poorly overall. Further, transformers were significantly inferior to models explicitly designed to encode the structural relations between words. Our results provide insight into the nature of sentence representation in the brain, highlighting the critical role of sentence structure. They also cast doubt on the claim that transformers represent sentence meaning similarly to the human brain.

---

Understanding how human language is processed and represented in the brain is a major scientific challenge. The past decade has seen a proliferation of work integrating theoretical approaches from linguistics and computer science with empirical data from neuroimaging studies in an effort to better understand how meaning is represented in the brain^1–7^. Most research has focused on evaluating vector-based models, in which the meaning of a word or phrase is represented as a vector of numbers. This approach forms the basis for large language models, which are neural networks based on the transformer architecture and trained to predict hidden tokens on very large corpora of natural text. Leading models such as GPT-4, Gemini, Llama, and Claude are highly versatile, capable of generating grammatical and relevant responses to a wide range of queries and instructions^8–10^. The extensive linguistic capabilities of these models, along with their ability to acquire language competence from naturalistic data, has generated significant interest in their potential value as cognitive models of language processing in humans^11–13^. Studies have consistently found statistically significant correlations between brain activity and various semantic models, with several finding that transformers better explain brain activity compared to static word embedding models^14–17^.

Most research comparing language models to brain activity has used stimuli that have not been selected to evaluate any specific linguistic hypothesis. While there are many benefits to utilising naturalistic stimuli in the study of language^18–21^, such stimuli have the disadvantage that they may not adequately sample the linguis-tic phenomena of most interest^19^, and do not control for variables crucial for contrasting the representations of different models^22^. A particular challenge is distinguishing whether language models are predictive of brain activity solely due to word-level (lexical) semantic information, or whether they also incorporate representations of sentence structure in a manner comparable to the brain. Direct comparison of static word embeddings with contextualised transformer embeddings is insufficient to resolve this issue, because contextualised embeddings also capture polysemy and other semantic phenomena not directly related to sentence structure. Another limitation of existing studies is that establishing that features extracted from large language models are predictive of brain activity does not necessarily provide much insight about what information these features encode or how such information is utilised by the brain^23–25^. A final limitation of existing studies is that encoding techniques are best suited to use with vector representations of language, making it difficult to conduct comparisons with graphbased or other approaches specialised for explicitly representing sentence structure.

Here, we present results from an fMRI study in which 30 participants read isolated sentences and answered simple questions about their meaning. We also collected a separate dataset of behavioural ratings of all pairwise comparisons of the same set of sentences. First, we developed a handcrafted set of sentences designed specifically to control for the confound of lexical similarity, allowing for clearer inferences about how sentence-level information is represented by the brain. Second, we conduct model comparison using representational similarity analysis (RSA), which involves comparing pairwise similarity scores for voxel activations and semantic models.

This technique extracts information about the patterns of similarity of model representations, thereby providing additional insight into the nature of brain semantic representations beyond voxelwise predictability. Furthermore, RSA facilitates comparison between dissimilar types of representations, thereby allowing us to compare a wider range of computational models, including both vector-based and graph-based models, than has been assessed by most previous research^26–30^.

## 1 Results

### 1.1 Stimuli and models

Our handcrafted sentences were carefully designed to reveal the role of sentence structure in semantic representation. Illustrative example sentences are shown in Figure 1a, along with the design matrix indicating the different types of sentence comparisons we considered. This matrix exhibits a block diagonal structure owing to the use of six distinct subsets of sentences each sharing a similar set of words. Within each of the six subsets, we begin with a base sentence such as ‘the cameraman brought the equipment to the director’, which we then systematically modified in various ways to create different combinations of lexical and compositional similarity, in order to dissociate these two aspects of meaning (see Table 1 for further details). We distinguish between ‘on-diagonal’ and ‘off-diagonal’ sentence pairs. On-diagonal sentence pairs (depicted in shades of blue) have sentence elements simply added or removed. By contrast, the off-diagonal sentence pairs (depicted in light green) have sentence elements interchanged to vary sentence meaning while keeping most of the constituent words the same. This approach builds on our previous work using behavioural data^31^, where we showed that such methods allow for effective dissociation of lexical similarity from overall similarity in sentence meaning. We explain the process for constructing the sentences in subsubsection 3.1.1. The primary objective of the present study is to analyse the brain representations of the block diagonal sentences extracted during an fMRI reading task, and compare these to representations derived from a variety of computational models of sentence meaning to determine which models best match brain representations.

**Table 1:** Explanation of the process of constructing sentences used in the study. Added or altered elements in the second sentence in each pair are italicised. The final two columns represent approximate relative similarities intended for each sentence pair type, though there will be variation due to the precise details of each sentence.

| Type | Example sentence pair | Lexical similarity | Overall semantic similarity |
| --- | --- | --- | --- |
| Same | The cameraman brought the equipment to the director.<br>The cameraman brought the *new* equipment to the director. | Highest | Highest |
| Modified | The cameraman brought the equipment to the director.<br>The cameraman *begrudgingly* brought the *awkward* equipment *up from the basement* to the director. | Moderate | High |
| Substituted | The cameraman brought the equipment to the director.<br>The cameraman *sold* the equipment to the director. | Moderate | Moderate |
| Modified & substituted | The cameraman *sold* the equipment to the director.<br>The cameraman *begrudgingly* brought the *awkward* equipment *up from the basement* to the director. | Low | Moderate |
| Swapped (simple case) | The cameraman brought the equipment to the director.<br>The *director* brought the *cameraman* to the *equipment*. | High | Moderate |
| Swapped (with substitution) | The cameraman brought the equipment to the director.<br>The *director* *sold* the *cameraman's equipment*. | Moderate | Low |
| Swapped (with modifiers) | The cameraman brought the equipment to the director.<br>The *renowned director* brought the *impatient camera-man* to the *equipment from his last project*. | Low | Low |
| Different | The cameraman brought the equipment to the director.<br>The psychologist warned the secretary about the patient. | Lowest | Lowest |

**Fig. 1:**
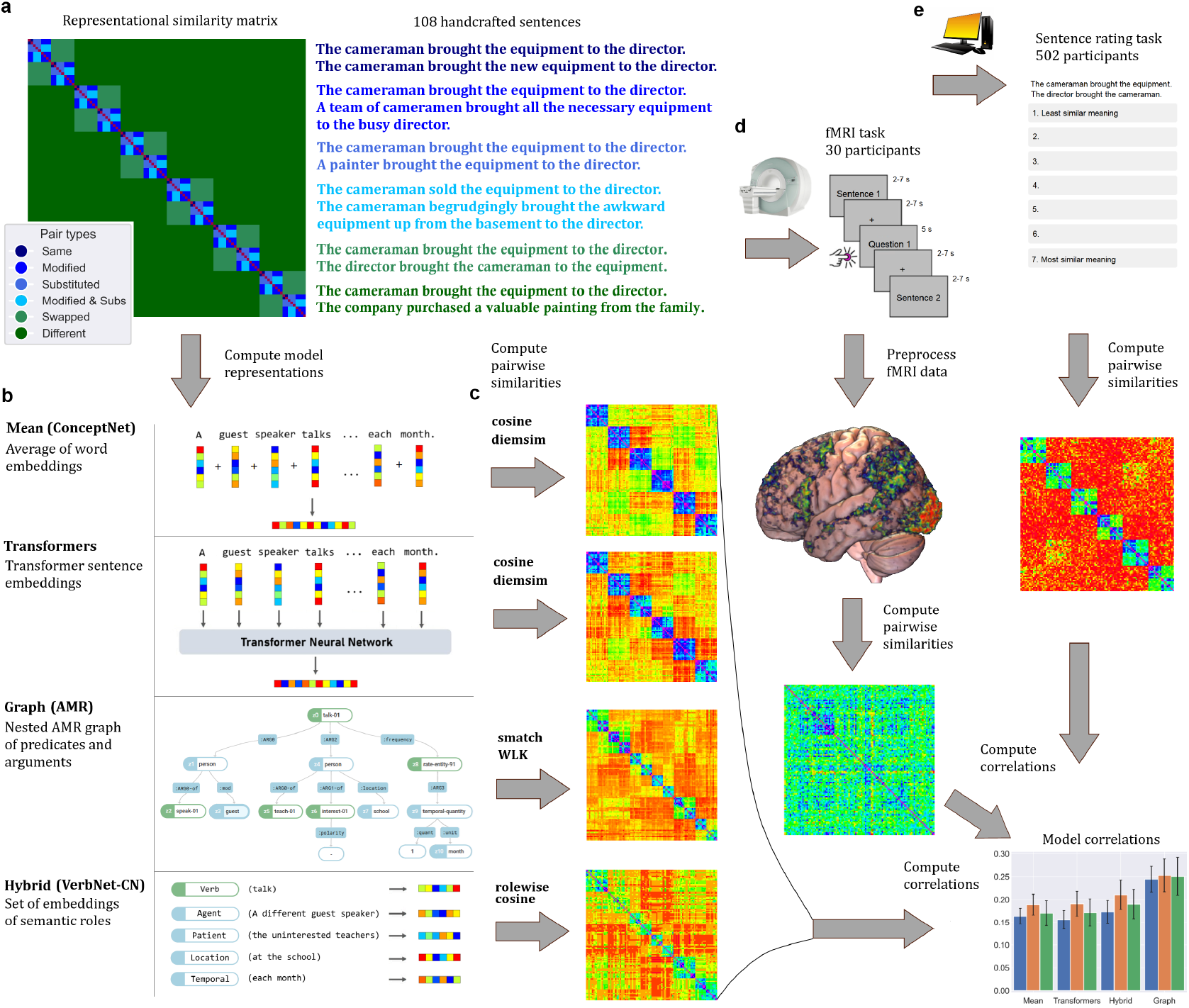
Summary of study methods for constructing stimuli, computing model representations, and collecting fMRI and behavioural data. **a)** We construct 108 handcrafted sentences, designed to enable systematic variation in sentence meaning while controlling for lexical similarity. Here we show the corresponding 108*×* 108 design matrix colour-coded with the type of each sentence pair. Sentence pairs in the six blocks along the diagonal are the primary pairs of interest in this study. **b)** All sentences were encoded using each of the four computational models of sentence meaning which we examine in this study. **c)** We then computed representational similarity matrices of the 108 stimuli for each of the four models. More similar sentence pairs are shown in blue, and less similar in red. **d)** Study pipeline for the fMRI experiment, in which participants were presented one sentence at a time for 2-7 seconds depending on sentence length. Multiple choice comprehension questions were interspersed randomly to assess attention. After scanning, data was processed and brain activity patterns were used to compute a neural representational similarity matrix for each participant. Correlations were then computed between the model and brain RSA matrices. **e)** Study pipeline for behavioural experiment, in which online participants were each shown 112 sentence pairs and asked to rate their semantic similarity. Ratings were averaged over participants to compute a similarity matrix. The correlation was then computed between the model and behavioural RSA matrices.

We next computed the representations for each sentence using a range of computational models. We analysed four distinct approaches to semantic representation. The first was a simple ‘Mean’ model, consisting of the element-wise averages of static word embeddings of each word in the sentence. Since this model ignores the position of words within a sentence as well as their grammatical role, it serves as a baseline incorporating only lexical information. The second class consists of embeddings extracted from various transformer neural networks. Results for the ‘Transformer’ model are calculated by computing correlations separately for five different transformer models and then taking a simple average of these correlations (details given in Methods subsection 3.1). Results for each individual transformer are presented in Figure S2. Both Mean and Transformer models are vector-based approaches, as they represent the meaning of a sentence with a vector of numbers^32^. By contrast, ‘Graph’ models are based on a nested graph formalism constructed in accordance with a semantic parsing paradigm. Here we selected Abstract Meaning Representation (AMR) as a widely-used exemplar of this approach to semantic representation^33^. Finally, we analysed a ‘Hybrid’ model called ‘VerbNet-CN’, which includes components from both vector-based and graphbased formalisms. Building on our previous work^31^, our VerbNet-CN model uses a semantic parser to tag each word based on its semantic role, and then constructs a separate vector embedding for each semantic role. All four models are summarised in Figure 1b.

Having constructed the model representations for our sentences, we next computed the similarities between all sentence pairs, using these data to construct RSA matrices for all four computational models. As shown in Figure 1c, the block diagonal structure corresponding to the six sentence subsets is clearly visible. Sentence pairs within these blocks have higher similarity owing to sharing many words in common, as per our design. More importantly, the RSA matrices also illustrate clear differences between how the four models represent sentences. In particular, the ‘swapped’ off-diagonal sentence pairs are accorded high similarities by the Mean-CN model, much lower similarities by the AMR-Smatch and VerbNet-CN models, and intermediate similarities by the Transformer models (OpenAI embeddings shown for illustration). These differences are consistent with our previous findings that transformers are less sensitive to changes in sentence structure than hybrid or graph models. Here we aim to test which pattern of representational similarities best matches data collected using neuroimaging during a sentence reading task. The full set of RSA matrices for all models is shown in Figure S1.

### 1.2 fMRI results

To evaluate how well each model describes sentence processing in the brain, we collected fMRI data from 30 participants while reading each of the 108 sentences. Our experimental pipeline is depicted in Figure 1d, with additional details given in section 3. We presented each sentence four times, with randomly interposed questions incorporated as an attention check. Voxel data were analysed using GLMsingle, an algorithm which fits a haemodynamic response function to each voxel and then estimates the response of that voxel to each stimulus. We selected a subset of voxels for further analysis based on their stability score, which is computed as the average correlation of voxel activity across repetitions of the same stimulus^2,34,35^. We analysed stable voxels within two regions of interest: the language network^36^, and the entire cortex less the primary visual cortex. Model fit was assessed using representational similarity analysis, with higher correlations indicating that the corresponding model represents the set of stimuli more similarly to the brain.

We performed representational similarity analysis in two different ways. In the simple-average approach, we computed the RSA correlation for each participant separately and then took the average. In contrast, the group-average approach involves first averaging the RSA matrix across participants, and then computing the RSA correlation for this group-averaged matrix^26,27,37^. In each case, we computed the Spearman partial correlation across all 5,778 sentence pairs and also across the 918 block diagonal sentence pairs, controlling for differences in sentence length. The full set of results for all 21 models tested is shown in Figure S2. Here we discuss results of the four models of main interest.

We first consider correlations computed using all sentence pairs, as shown in Figure 2a. In language network voxels, all models show positive correlations, with relatively small differences between models. For the simple-average method, the differences in correlation were not significant when comparing the Mean and Transformer models (Δ*ρ* = 0.001, *t* = 0.686, *p* = 0.4981), and only marginally significant (after multiple comparison correction) for the VerbNet-CN and Transformer models (Δ*ρ* = 0.009, *t* = 2.720, *p* = 0.0109). However, the AMR-Smatch model had a significantly higher correlation compared to the VerbNet-CN model (Δ*ρ* = 0.043, *t* = 7.393, *p* < 0.0001). Similar results were found using the group-average method (shown in Figure 2b), but with higher absolute values. The fact that all models show positive correlations when evaluating all sentence pairs is unsurprising, since most sentences can be differentiated from one another using purely lexical differences, which all models are sensitive to.

**Fig. 2:**
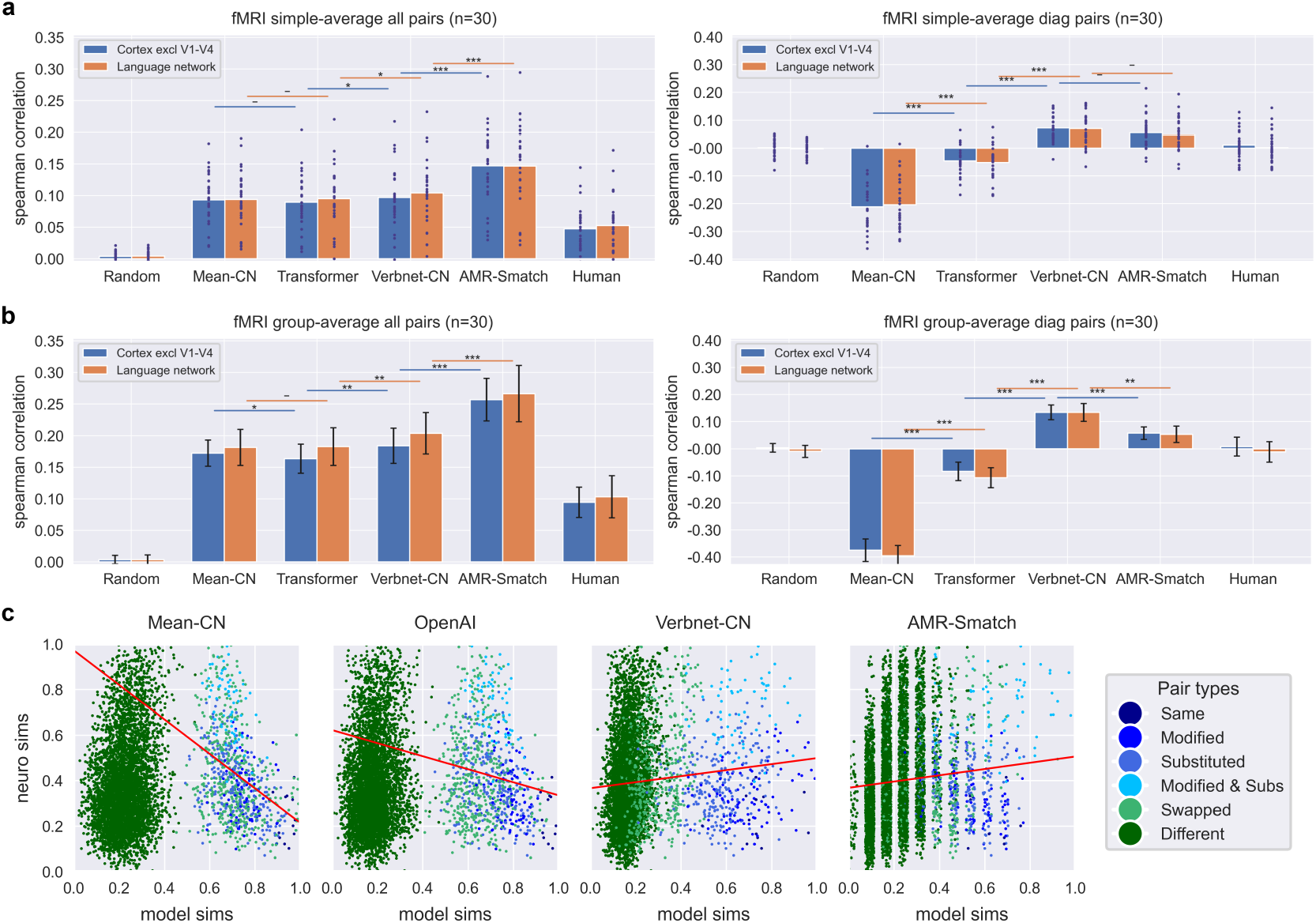
Model correlations with brain activity for all sentence pairs and the block-diagonal subset of sentence pairs. Partial correlations between RSA matrices of five computational models (Random, Mean, Transformer, VerbNet-CN, and AMR-Smatch) and the brain RSA matrix, controlling for differences in sentence length. ‘Human’ refers to behavioural ratings. Blue bars show correlations computed over all stable voxels (excluding visual regions V1-V4), while orange bars show correlations for stable voxels in the language network. Notation for statistical significance: * for p<0.05, ** for p<0.01, and *** for p<0.001, with Bonferroni correction for three independent comparisons. **a)** Partial correlations for each individual participant shown as blue dots, with the simple average over individual correlations shown as a bar. **b)** Partial correlations computed using the group-averaged RSA matrix. Error bars show 95% confidence intervals calculated by bootstrap resampling over participants. **c)** Scatterplots showing the relationship between model similarities (horizontal axis) and group-average neural similarities (vertical axis) for all four computational models. Each dot corresponds to a single pairwise similarity, scaled to between 0 and 1 for visualisation. While all sentence pairs are shown for comparison, regression lines (red) are computed over the block diagonal pairs only.

We now consider correlations computed using only the block diagonal sentence pairs, which are designed to be more difficult for computational models to distinguish owing to high lexical similarity. Here our results are noticeably different. For the simple-average of voxels within the language network, we found a correlation of −0.204 for the Mean-CN model. This comparatively large negative correlation indicates that brain representations of sentences differ systematically from representations constructed considering only lexical similarity, providing evidence that brain representations of sentences are highly sensitive to sentence structure. The Transformer model achieves a correlation of −0.045, which is significantly higher than the Mean-CN model (Δ*ρ* = 0.159, *t* = 14.287, *p* < 0.0001), though the negative sign indicates that transformers still poorly match brain similarities, and indeed more closely resemble the static word embeddings that ignore sentence structure. The VerbNet-CN model achieves the highest correlation of 0.070, much larger than the Transformer model (Δ*ρ* = 0.115, *t* = 8.150, *p* < 0.0001). The AMR-Smatch model shows similar results to the VerbNet-CN model, with a correlation of 0.047 (Δ*ρ* = −0.023, *t* = −1.783, *p* = 0.0851). The correlation with human ratings is very close to zero, placing it between the Transformer and VerbNet-CN models. We consider this surprising result in greater detail in section 2. Results were very similar using the group-average method, though generally correlations had higher absolute values.

The results for Mean, Transformer, and VerbNet-CN models were all consistent with our preregistered predictions based on previous work with a separate behavioural dataset^31^, though we did not make a prediction for the AMR-Smatch model. In all cases, results are very similar whether computed over the entire cortex (excluding V1–V4) or focusing just on the language network. Results are similar when using the DIEM similarity metric instead of cosine similarity, though with somewhat lower correlations for certain transformer models (see Figures S12 and S13).

To better understand the origin of such large differences in correlations, we plotted neural similarities against the similarities derived from all four computational models (see Figure 2c). For both the Mean and Transformer models, the blue ‘modified’ and ‘substituted’ sentence pairs are accorded comparable similarities to the light green ‘swapped’ sentence pairs. By contrast, the VerbNet-CN and AMR-Smatch models generally accord ‘swapped’ sentence pairs as having distinctly lower similarity than ‘substituted’ and ‘modified’ sentence pairs. This is easiest to see on the VerbNet-CN subplot of Figure 2c, where the ‘swapped’ sentence pairs are noticeably to the left of the ‘modified’ and ‘substituted’ sentence pairs. Such a difference indicates that the VerbNet-CN and AMR-Smatch models have a greater ability to discriminate sentence pairs that are lexically similar but structurally different (due to interchanged semantic roles). This leads to sentence similarities which are in better accord with brain similarity data, and thereby drives the positive RSA correlations. These results indicate that when keeping lexical similarity roughly constant, as is the case for the block diagonal sentence pairs, brain similarity patterns are best explained by models that explicitly represent sentence structural elements, namely the VerbNet-CN and AMR-Smatch models. The Mean-CN model, which completely ignores such structure, explains brain representations the worst, with Transformer models doing better than the Mean but still poorly overall.

We next compared representations across different brain regions. In addition to the language network and visual cortex (V1–V4), we also considered several regions previously demonstrated to show activity in response to language stimuli, namely the dorsomedial prefrontal cortex, the dorsolateral prefrontal cortex, the posterior cingulate cortex, and the precuneus. The primary somatosensory cortex (S1) is also included as a comparison of a brain region expected to show little response to linguistic stimuli. As shown in Figure 3a, the RSA matrices for most of these regions show a very robust grid-like pattern not explained by the type of sentence pair in the design matrix. This effect is not explained by differences in sentence length, as the RSA matrices already control for this variable (shown on the right of Figure 3a). Upon further investigation, we identified the grid-like pattern as resulting from consistently high brain similarity of sentence pairs in which both sentences are relatively long, as measured by the number of characters. This is evident by visual comparison with the ‘minimum length’ RSA matrix on the right of Figure 3b, which shows the shortest length of the two sentences in each pair. In Figure S4, we show that our main results are qualitatively similar when additionally controlling for the ‘long sentences effect’. After regressing out this effect using the minimum sentence length for each sentence pair (Figure 3b), we recovered a block diagonal structure comparable to the original design matrix shown in Figure 1a, most clearly visible in the language network. As an additional check, we computed correlations controlling for the fMRI similarities computed over the visual cortex (V1-V4) averaged over all participants. Even with this very strict control of visual similarity, we still observe the same pattern of similarities across the four models (see Figure S8).

**Fig. 3:**
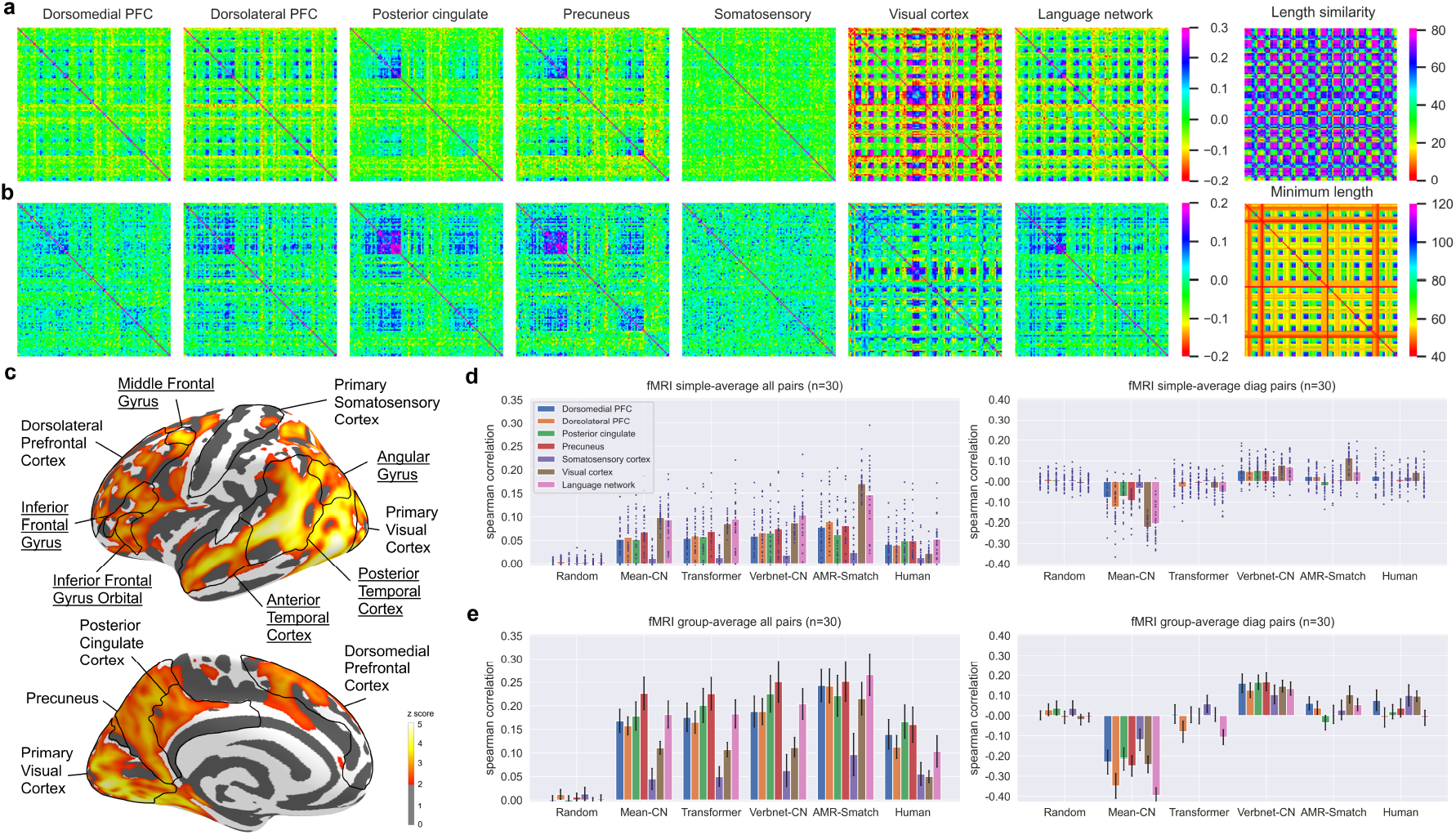
Comparison of sentence representations and model correlations across brain regions. **a)** RSA matrices for various cortical regions, computed controlling for differences in sentence length. **b)** RSA matrices for various cortical regions, computed controlling for differences in sentence length and minimum sentence length. **c)** Searchlight RSA for the VerbNet-CN model using 8mm radius showing cortical regions of interest, with those part of the language network underlined. RSA correlations are thresholded at z=2. **d)** Partial correlations controlling for differences in sentence length by cortical region, with each individual participant shown as blue dots, and the simple average over individual correlations shown as a bar. **e)** Partial correlations controlling for differences in sentence length computed using the group-averaged RSA matrix, shown by cortical region. Error bars show 95% confidence intervals calculated by bootstrap resampling over participants.

We also conducted an analysis of RSA correlations for each layer of the Llama 3 transformer model. We chose this for analysis as a larger, more recent architecture with a large number of layers. As shown in Figure 4, layers 0 and 1 had large negative correlations more similar to the Mean-CN model, while layers 2 and 3 had slightly positive correlations closer to that of the VerbNet-CN model. Layers 4 and on had more moderate negative correlations, with a slight downward trend over later layers. This pattern was largely similar for both the set of all pairwise comparisons and the set of block diagonal comparison pairs, though in the latter case correlations remained essentially constant from around layer 4 onwards. The corresponding RSA matrices (see Figure 4c) show clear differences in representation across layers, with the earlier layers in particular showing evidence of representations dominated by the effects of sentence length and visual similarity (compare with RSA matrices for Lengthsim, Length-min, and Visual in S1). This is dramatically evident when controlling for visual similarities, as this results in correlations over all sentence pairs falling significantly below that of VerbNet-CN, with the highest correlations now observed around layer 28 instead of layer 2 (see Figure S11). We found only modest differences across layers of the AMRBart and ERNIE transformers (see Figures S16 and S17).

**Fig. 4:**
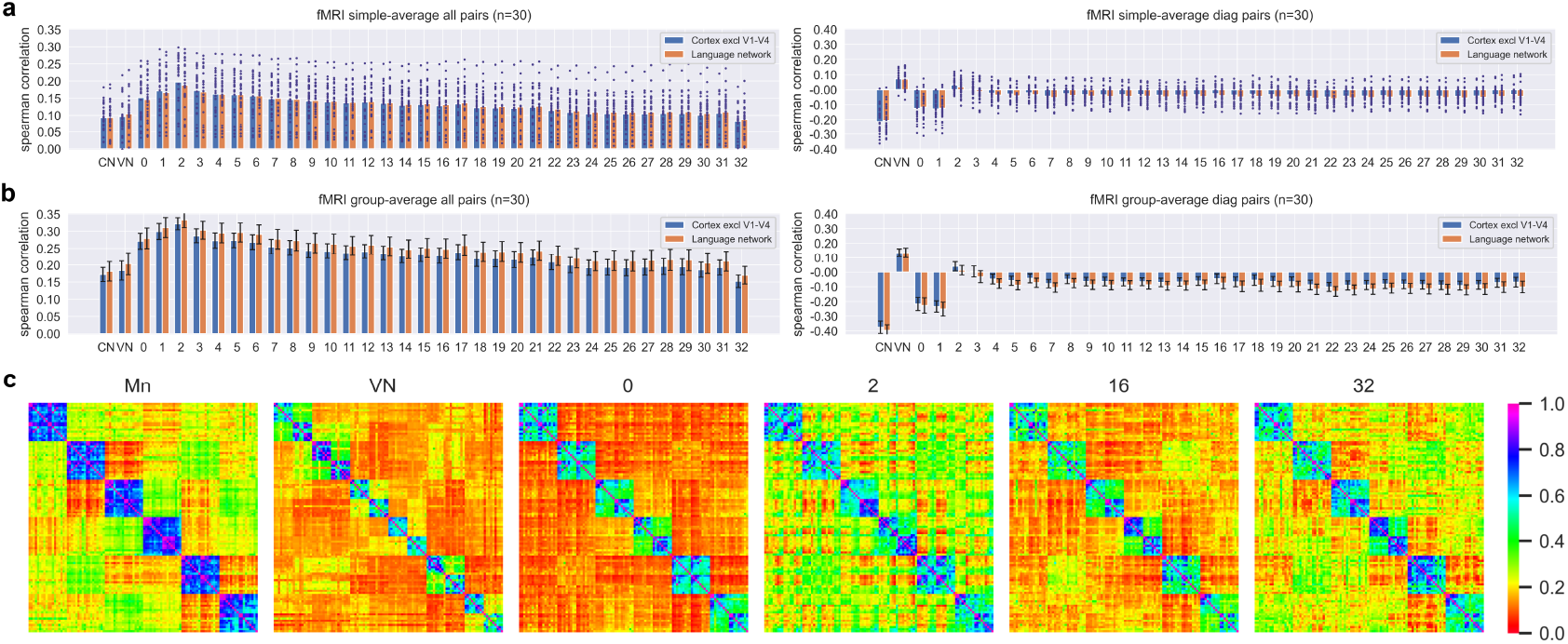
Average correlations between RSA matrices of each layer of Llama 3 and brain RSA matrix of each participant. Mean-CN (CN) and VerbNet-CN hybrid (VN) models are also shown for comparison. **a)** Partial correlations for each individual participant shown as blue dots, with the simple average over individual correlations shown as a bar. **b)** Partial correlations computed using the group-averaged RSA matrix. Error bars show 95% confidence intervals calculated by bootstrap resampling over participants. **c)** RSA matrices for the Mean-CN model along with selected layers of the Llama 3 model, computed controlling for differences in sentence length.

To more clearly visualise the location of the brain regions responsible for encoding sentence information in common with the computational models, we conducted an RSA-searchlight analysis. This involves computing the RSA correlation between each model and the voxel activations within an 8mm sphere surrounding each voxel within a cortical mask. The results (see Figure 3c) show significant correlations throughout the language network, including regions of the temporal lobe, the angular gyrus, and the frontal lobe. Significant correlations are also evident in the posterior cingulate cortex, precuneus, and the visual cortex, with sporadic pockets throughout the dorsolateral and dorsomedial frontal cortical regions. In Figure 3d-e we show the correlations for each model in each cortical region. We observe low correlations for the somatosensory cortex, generally high correlations for the language network, and intermediate correlations for all other regions. For block diagonal sentence pairs, the VerbNet-CN model has similar correlations across all regions, while the AMR-Smatch model has the highest correlation in the visual cortex, but still positive correlations in the language network. We find similar results when additionally controlling for minimum sentence length, as shown in Figure S5.

We also performed an analysis comparing the representation of each subregion of the language network, the locations of which are depicted in Figure 5a. We found a similar overall pattern of results within all subregions, with consistently positive correlations for the entire set of pairwise comparisons. The magnitude of the correlations varied across subregions, with the highest values observed for the anterior and posterior temporal lobe, and lower values for all frontal regions (see Figure 5b-c). For the set of block diagonal sentence pairs, all subregions showed the same pattern as our main results, with a negative correlation for the mean model, modest negative correlations for transformer models, and positive correlations for the hybrid model. These findings support previous results indicating that all subregions of the language network are sensitive to lexical, syntactic, and compositional aspects of language, without any obvious specialisation across subregions^38,39^. We find little difference when additionally controlling for minimum sentence length, as shown in S6.

**Fig. 5:**
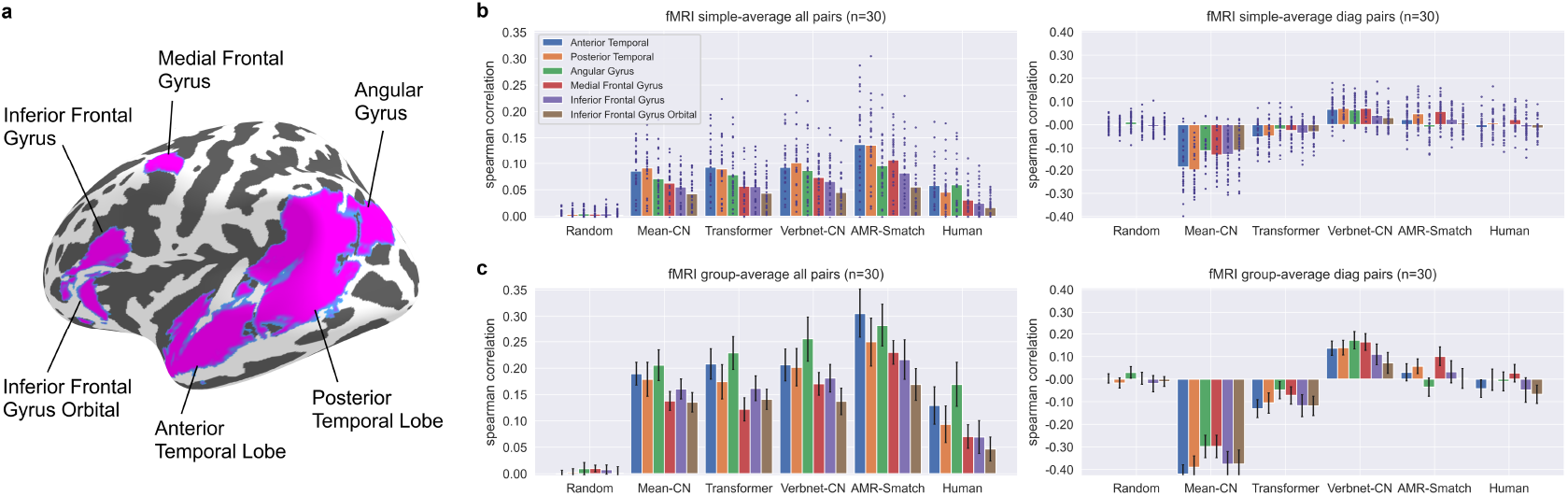
Comparison of model correlations across subregions of the language network. **a)** Regions within the language network. **b)** Partial correlations controlling for differences in sentence length by brain region, with each individual participant shown as blue dots, and the simple average over individual correlations shown as a bar. **c)** Partial correlations controlling for differences in sentence length computed using the group-averaged RSA matrix, shown by language network region. Error bars show 95% confidence intervals calculated by bootstrap resampling over participants.

### 1.3 Behavioural results

To supplement our neuroimaging data, we also collected a set of behavioural data consisting of semantic similarity judgements. As illustrated in Figure 1e, we recruited 502 participants using an online platform, each of whom was presented with a set of 112 sentence pairs selected randomly from all 5,778 unique sentence pairs. Participants were asked to rate each sentence pair for semantic similarity on a scale of 1-7. Ratings were averaged over participants and scaled to between 0 and 1 for comparison with model similarities. The normalised human sentence similarity ratings ranged from 0 to 0.962, with mean=0.484 and SD=0.171 for block diagonal sentence pairs, and mean=0.072 and SD=0.071 for all other sentence pairs. The average standard deviation of similarity scores for each sentence pair computed across participants was equal to 0.244 for block diagonal sentence pairs and 0.106 for all other pairs. This is comparable to the 0.19 adjusted average standard deviation of the SICK sentence similarity dataset^40^, and 0.216 for the STS3k dataset^31^. The split-half reliability with the Spearman-Brown correction was 0.938 for the entire dataset, 0.954 for the block diagonal sentence pairs, and 0.715 for all other pairs, indicating high levels of agreement between participants.

We evaluated the fit between behavioural data and each computational model in the same manner as for the fMRI data. For the full set of sentence pairs (Figure 6a left), the Mean and Transformer models performed best with correlations of 0.510 and 0.568 respectively (Δ*ρ* = 0.049, *t* = 11.327, *p* < 0.0001). The VerbNet-CN model had a lower correlation relative to the Transformer (Δ*ρ* = −0.093, *t* = −16.432, *p* < 0.0001), and the AMR-Smatch model the lowest of all (Δ*ρ* = −0.044, *t* = −6.306, *p* < 0.0001). This pattern was reversed in the case of the block diagonal sentence pairs (Figure 6a right), with the Mean-CN model having by far the lowest correlation of 0.437. Transformer had a much higher correlation of 0.639 (Δ*ρ* = 0.188, *t* = 22.449, *p* < 0.0001), as did the VerbNet-CN model with a correlation of 0.698 (Δ*ρ* = 0.045, *t* = 3.765, *p* = 0.0001). The AMR-Smatch model had an intermediate correlation of 0.533, lower than the VerbNet-CN model (Δ*ρ* = −0.145, *t* = −12.371, *p* < 0.0001). This pattern of results is comparable to that we observed for our fMRI data, though with much higher correlations across all models owing to the much reduced noise in behavioural ratings compared to fMRI voxel data.

**Fig. 6:**
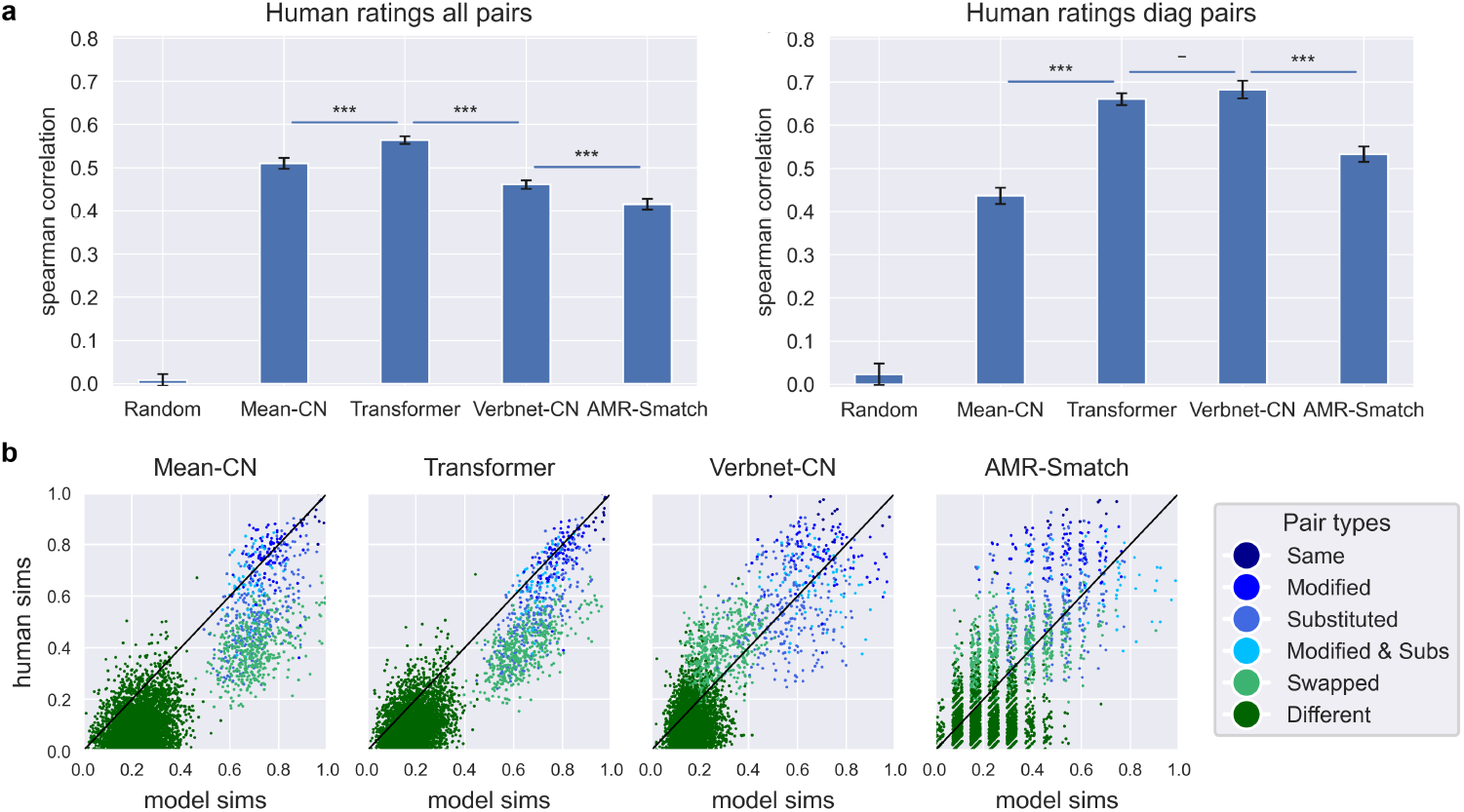
Behavioural ratings of sentence similarity show similar results to fMRI results, but with higher absolute correlations. **a)** (Left) Average correlations between RSA matrices of four computational models and human-rated similarities using all sentence pairs. (Right) Average correlations between RSA matrices of four computational models and human-rated similarities using only diagonal sentence pairs. **b)** Scatterplots showing the relationship between model similarities (horizontal axis) and human rated similarities (vertical axis) for all four computational models. Each dot corresponds to a single pairwise similarity, scaled to between 0 and 1 for visualisation. The 45-degree line (black) shows a hypothetical line of perfect fit between model and human similarities.

As a supplementary analysis not in our original pre-registration, we also asked GPT-4 to directly provide ratings for the semantic similarity of each pair of sentences. We found that over both the full set of sentences and the block diagonal subset, these ratings achieved correlations higher than any other method, with values of 0.616 and 0.838 respectively. Correlations for all computational models are shown in Figure S3.

As before, we show scatterplots of the human ratings plotted against model similarities (Figure 6b). While all four models broadly follow the ordering of human ratings along the 45-degree line, both the Mean and OpenAI transformer models place the ‘swapped’ sentence pairs below the line, meaning that these sentence pairs are accorded higher similarity ratings by the models than by humans. By contrast, the VerbNet-CN and AMR-Smatch models place ‘swapped’ sentence pairs above the 45-degree line, meaning that they accord these sentence pairs lower similarities than humans do. These results indicate that for this set of stimuli, the Mean and OpenAI transformer models are less sensitive to variations in sentence structure than human raters, while the VerbNet-CN and AMR-Smatch models are slightly more sensitive to such structure than the human raters.

## 2 Discussion

In this paper we present, to our knowledge, the first fMRI evaluation of models of sentence representation that utilises stimuli specifically designed to distinguish the effects of lexical semantics from sentence structure. We also present the first quantitative comparison of static word embeddings, transformer neural networks, semantic parsing graphs, and hybrid representational models all under a unified framework. In our neuroimaging experiment we found that over the block diagonal sentence pairs (the subset of sentence pairs designed to test for sensitivity to sentence structure), considering voxels in the language network, the Mean-CN model had a strong negative correlation, the Transformer model a smaller negative correlation, and the VerbNet-CN model a modest positive correlation. We found similar (though less pronounced) differences in our behavioural experiment. These findings provide two major contributions to our knowledge of sentence representation in the brain. First, we show that controlling for lexical similarity illuminates the brain’s sensitivity to sentence structure in a way that is not evident when the lexical confound is present. Second, the success of our VerbNet-CN model provides novel insight into how sentence structure is represented in the brain, indicating the importance of semantic roles and highlighting limitations of representations derived from transformer models.

### 2.1 Comparison with previous work

Previous studies analysing sentence processing in the brain have used a variety of controlled stimuli to isolate the mechanisms of semantic composition. One method involves randomly shuffling the order of words within a sentence, thereby preserving lexical semantics while varying overall sentence meaning^41,42^. A second method involves constructing ‘jabberwocky’ sentences, which involve nonsensical words placed in grammatically well-formed sentences^38,43,44^. These stimuli are designed to control for syntactic structure or sentence form while manipulating sentence meaning. In both cases, the objective is typically to use jabberwocky or shuffled sentences as a control condition in which composition is prevented, thereby providing a baseline for sentences in which composition occurs^45^. Our study differs from these approaches in that we aim to preserve, rather than prevent composition. Instead, we control for lexical similarity while constructing semantically meaningful sentences with differing meanings.

Another approach that has seen widespread use is the presentation of minimal sentence pairs that differ only in one specified aspect, for example interchanging subject and object in a sentence^46–49^, or altering adjective-noun phrases to influence composition^50–53^. Our approach is an extension of these techniques that utilises more naturalistic and complex sentences, designed to facilitate comparison of a wider range of structural manipulations (see Table 1). By more completely characterising the representational structure of various computational models in response to various semantic contrasts, we are able to more comprehensively evaluate their adequacy as models of semantic processing in the brain.

### 2.2 Transformer models

Our results indicate that transformer representations do not adequately incorporate sentence structure in a brainlike manner. While most models perform well when evaluated over the full set of sentence pairs, when evaluated against the block diagonal pairs only, transformers are insufficiently sensitive to ‘swapping’ of semantic roles (see Figure 6 and Figure 2), ranking such sentence pairs as more similar than do human participants (in the behavioural data) or brain representations (in the fMRI data). This effect was very robust, with negative correlations observed for all transformers and static word embeddings we studied aside from DefSent (see Figures S2 and S7). These negative correlations result from our experimental design, and arise for models which accord high similarities to sentence pairs that are lexically similar, but differ in meaning due to interchanging of semantic roles (see Table 1 for examples). These semantic differences lead to relatively dissimilar brain representations, in contrast to the high similarities of the model representations. The resulting negative correlations therefore indicate that transformer representations are much less sensitive than humans to the effect that semantic roles have on overall sentence meaning.

Similar results were found with the DIEM similarity metric, with negative correlations observed for block diagonal pairs for all transformer and static word embedding models except DefSent and GloVe (see Figures S13 and S14). When visual similarities were controlled for, all transformer and static word embedding correlations fall dramatically, with even DefSent and GloVe becoming negative, indicating that they may be significantly influenced by factors correlated with visual features, such as sentence length or superficial form (see Figures S9, S10, and S15). By contrast, correlations for the AMR-Smatch and VerbNet-CN models are much less affected after introducing controls for visual similarity, with the VerbNet-CN model in particular maintaining a positive correlation over block diagonal sentence pairs for every combination of similarity metric and control condition.

Several previous studies have found that voxelwise encoding models trained using features extracted from transformers are able to better predict brain activity than static word embedding models which ignore sentence structure^15–17,54^. However, interpreting these findings is difficult because there is no established method for determining which model features drive these correlations^55^. Indeed, some studies have found that even features from untrained transformers can achieve high voxelwise correlations^15,56^, casting doubt on whether the transformer features which drive brain correlations are linguistically relevant. Similarly, other studies using shuffled sentences to remove information about sentence structure have found this results in only modest reductions in voxelwise correlations^57,58^. An analyses which better controlled for various confounds found that most variance explainable by transformers was accounted for by static word embeddings and word rate^59^. Our results complement these findings, showing that in cases where sentence structure is critical, transformer representations are insufficiently sensitive to structural aspects of sentence meaning. In cases where transformers have been found to have an advantage, this may be due to their greater ability to contextualise polysemous word meanings based on the presence of other words, rather than their ability to represent sentence structure.

We emphasise that our results do not show that transformers fail to represent syntactic or semantic role information. Indeed, large language models show clear capabilities of correctly interpreting sentence structure^60^, and probing studies have found that transformers represent information about syntax and word order^61,62^. This is consistent with our finding that directly prompting GPT-4 to rate sentence similarity yields very high correlations with human judgements (see Figure S3). Nonetheless, the fact that transformers can encode and utilise structural information to perform linguistic tasks does not mean that they effectively utilise this information to construct a brain-like representation of sentence meaning. Our results indicate that despite the linguistic competencies of transformers, when controlling for lexical similarity, transformers do not combine syntactic and semantic information into an integrated sentence representation in a manner analogous to the human brain. Another problem with using transformers as models of semantic representation is that they are largely ‘black box’ models whose representations are often difficult to understand. Graph-based and hybrid models, whose semantic representations are much more transparent and interpretable, can thus play an important role in increasing our understanding of how semantic information is represented by large language models, and to what extent such representations differ from those formed in the human brain.

### 2.3 Graph and hybrid models

Our results for the graph-based models were rather mixed. We found that purely syntactic models based on constituency parses (the Benepar and CFG models) have low correlations with brain activity (see Figure S2. Examining the corresponding RSA matrices (see Figure S1), this seems to be due to such models being overly sensitive to syntactic form, and relatively insensitive to which words are assigned to different nodes within the syntactic tree. This can be seen in the RSA matrices in the four blue squares within each of the six block diagonal squares, which indicates that the ‘swapped’ sentences are not adequately distinguished from ‘same’ sentences (compare with Figure 1a). Comparison with the Length-sim and Length-min RSA matrices also indicates that the edit-distance similarity metrics are strongly affected by sentence length. The AMR-WLK model shows a similar RSA pattern to the Benepar and CFG models, which may account for its low brain and behavioural correlations. Interestingly, the AMR-Smatch model has relatively high brain correlations, despite differing from AMR-WLK only in the similarity metric used. We speculate this could be explained by the fact that Smatch similarity is based on the number of node triples shared between two graphs, which could be more effective at encoding semantic roles than the more complex node-embedding method used in the WWLK metric (see subsection 3.1 for further details). These findings highlight the importance of carefully evaluating graph similarity metrics and identifying which are most appropriate for comparisons of semantic similarity. Several previous studies have likewise emphasised the limitations of existing metrics and the need to explore alternatives^63–65^.

The hybrid VerbNet-CN model achieves the highest brain correlations for on-diagonal sentence pairs of all models tested, and comparable behavioural correlations to leading transformer models. We believe this is due to this model being designed specifically to be highly sensitive to semantic roles, which is the major point of differentiation from most other models. Indeed, for the behavioural data we find that the hybrid model is actually more sensitive to these ‘swapped’ sentences compared to human participants, who rate their similarity in between that of the VerbNet-CN and Transformer models (see Figure 6). Interestingly, the second hybrid model we analysed, AMR-CN, shows low brain correlations (see Figure S2). We speculate this is likely due to the crude method in which AMR-CN extracts semantic roles from the uppermost layer of the AMR graph of each sentence, in contrast to the VerbNet-CN model which uses GPT-4 to identify semantic roles directly. Indeed, this difference is why we predicted that VerbNet-CN would perform best in our preregistration. We also wish to emphasise that our core message is not the superiority of the Verbnet-CN model per se, but that the better performance of the VerbNet-CN model over block-diagonal sentence pairs highlights that static word embeddings and transformer representations are insufficiently sensitive to semantic roles. These results highlight the value of hybrid approaches designed to appropriately balance sensitivity to lexical, syntactic, and compositional information in representing semantic information at the sentence level, while also indicating that details of how semantic features are extracted are critical for constructing brain-like sentence representations.

### 2.4 Neuroscience of semantics

Our neuroimaging results show that linguistic information is represented across large parts of the cortex beyond the language network, including the default mode network that has been implicated in semantic processing in previous studies^66–68^. This supports previous studies which have found that processing of lexical semantics is intermingled with syntactic and structural processing^38,69^. One interesting supplementary finding is that the temporal regions of the language network tended to show somewhat larger effects (i.e. more negative Mean-CN correlations and higher VerbNet-CN correlations over the block diagonal pairs) compared to frontal regions of the network (see Figure 5). This aligns with several previous studies which have similarly found regions of the temporal lobe, especially the superior temporal sulcus, to play a prominent role in compositional and sentence processing^30,47,70^.

We also found a robust ‘minimum sentence length’ effect, whereby long sentences elicit very similar brain activity regardless of their lexical content or overall meaning (see Figure 3a-b). This effect is specific to long sentences, and does not arise for pairs consisting of short or medium-length sentences. Though we are not aware of this result having been reported using RSA, previous studies using other methodologies have found that activation of the language network increases with sentence length^38,71–73^. The cause of this effect is unclear. It may partly be explained by visual similarity of long sentences, however we observe no similar effect that might be expected for the visual similarity of short or medium sentences. Furthermore, the minimum length effect is also evident in many brain regions outside the visual cortex, including the language network and various frontal regions Figure 3. We speculate that the effect may be driven by multiple causes, including increased cognitive processing or memory load for processing longer sentences, greater depth of processing elicited by semantically richer stimuli^74^, or additional processing required for compositional combination of a larger number of sentence components. It is also possible that the structural similarity of longer sentences in our study, which all contain a similar set of semantic roles, results in similar brain representations even when the sentences do not have similar overall meanings. If so, this would indicate that extracting semantic features is important for brain processing of sentences even aside from lexical similarity. Further research will be required to disentangle the relative impacts of these distinct processes.

### 2.5 Limitations

Our study has several limitations. First, we found a surprisingly low correlation between behavioural ratings and brain activations (see Figure 2). This may be partly explained by differences in task structure. In the behavioural experiment, participants viewed many pairs of related sentences, and were explicitly asked to pay attention to differences in the words of each sentence. Conversely, in the fMRI task participants (who were not the same as the behavioural task participants) read one sentence at a time without an explicit comparison. In addition, we suspect that presentation of so many sentence pairs with highly similar structures may have biased the way in which participants rated sentence similarity. Modifications to the behavioural task to mitigate these aspects may reduce the divergence between behavioural and brain findings.

Second, our stimulus set consists of a relatively small selection of sentences, which follow broadly similar structure. Our aim in this study was to disentangle the effects of lexical similarity from structural similarity in realistic sentences, and as such we did not attempt to compile a representative sample of sentences from natural dialogue. In future work we hope to investigate the extent to which our results generalise to more complex and varied types of sentences.

Third, we analysed brain representations of sentence meaning over a single contiguous 3s interval. This is a substantial simplification of sentence processing, which takes place dynamically over time as words are successively integrated to form progressively more complex and structured representations^22,38,75–77^. While our approach is an important contribution, and builds upon previous studies comparing syntactic parse trees with brain data^71,78,79^, additional work is needed to link model representations with the dynamic cascade of brain activity during sentence processing.

### 2.6 Conclusion

Our results provide important new insights about how sentence structure is represented in the brain. The simple Mean-CN model, which ignores sentence structure, was a very poor match to brain activity when evaluated against the block diagonal subset of sentences (the sentence pairs designed to be difficult for models which do not represent sentence structure). While transformers were a much better match to brain activity than the Mean-CN model, correlations were still negative, indicating that transformer representations were still a poor match to brain representations. In line with our preregistered prediction, we found that the VerbNet-CN model best matched brain representations, thereby providing evidence that the brain incorporates structured information from semantic roles when representing sentence meaning. Even though transformer models clearly do encode semantic role information, the fact that their hidden layers fail to show the similarity structure of human BOLD responses indicates that transformers do not represent this information in the same manner as the human brain. Our results therefore highlight the important lesson that human-like linguistic behaviour does not entail human-like linguistic representations.

## Acknowledgements

We would like to thank the staff at the Melbourne Brain Centre Imaging Unit for their assistance with collecting the fMRI scans.

## Funding

This research was supported by a University of Melbourne Graduate Research Scholarship from the Faculty of Business and Economics (Fodor).

## Author contributions

Conceptualisation: J.F.; Methodology: J.F., C.M., S.S.; Investigation: J.F.; Formal analysis: J.F.; Visualisation: J.F.; Supervision: C.M., S.S.; Writing – original draft: J.F.; Writing – review editing: J.F., C.M., S.S.

## Competing interests

The authors declare that they have no competing interests.

## Data and materials availability

Our fMRI dataset is available for download on Open-Neuro: https://openneuro.org/datasets/ds007393. All code is available via github: https://github.com/Fods12/sentence_meaning_in_the_brain.

## 3 Methods

### 3.1 Stimuli and computational models

#### 3.1.1 Sentence stimuli

A set of 108 sentences was handcrafted specifically for this study. Our aim was to develop a set of sentence pairs which systematically tested different combinations of lexical similarity and overall semantic similarity. This allows for better model discrimination by ensuring that only models sensitive to sentence structure are able to accurately differentiate sentence meaning, reducing the confound of lexical similarity.

The process by which sentences were constructed is summarised in Table 1. All sentences consisted of a single clause written in the active voice describing a specific event. Pronouns, proper nouns, and subordinate clauses were excluded for simplicity and to limit sources of syntactic variation. Sentences were produced by constructing systematic variations of an initial ‘base’ sentence by altering elements such as the subject, verb, and object, or adding modifiers like adjectives, location, or time. In an effort to explore different combinations of lexical and overall sentence similarity, several different categories of altered sentences were constructed. A small number of ‘same’ sentences were constructed by adding a single adjective with only minimal effect on sentence meaning, for example ‘the equipment’ becomes ‘the new equipment’. ‘Modified’ sentences were constructed by adding two or three modifier elements such as location, manner, or time when the event occurred, under the hypothesis that adding these modifier terms would reduce lexical similarity but have only a small effect on overall sentence meaning. ‘Substituted’ sentences were designed to investigate the effect of altering key sentence elements, such as changing the subject, object, or verb of the sentence.

Critical to the study design was construction of ‘swapped’ sentences, in which one or more pairs of words interchanged roles in the sentence, thereby ensuring that lexical similarity is high while similarity in overall sentence meaning is low. For example, if in the initial sentence the subject is ‘the cameraman’, the direct object ‘the equipment’, and the indirect object ‘the director’, then in the interchanged sentence the subject is now ‘the director’, the direct object is ‘the cameraman’, and the indirect object is ‘the equipment’. As with the ‘base’ sentence, the swapped sentences were also systematically varied through substitutions and addition of modifiers. The aim of this procedure was to develop a set of sentence pairs with gradations of similarity while approximately controlling for lexical similarity. Differences in meaning in these sets of sentences are therefore mostly attributable to sentence structure and semantic roles, not simply use of different words. The complete set of stimuli are provided in Supplementary Information subsubsection 1.2.1.

Using the methods described above, six distinct subsets each consisting of 18 related sentences were developed. This resulted in 5,778 pairwise comparisons across all sentences, of which 4,860 were ‘different’ sentence pairs and 918 were block diagonal sentence pairs of primary interest in this study. The RSA design matrix for all 108 sentences is shown in Figure 1a.

In our study preregistration (see https://osf.io/jme7x), we predicted that over the block diagonal set of sentence pairs, the VerbNet-CN model would have a higher correlation with brain representations than the average over the five specified transformers, which in turn would have a higher correlation than the Mean-CN model. We did not make predictions for any other models.

#### 3.1.2 Word embedding models

In this study we compared four different approaches for representing sentence meaning. The baseline for all comparisons was the Mean-CN model in which sentence embeddings are constructed by elementwise averaging of word embeddings. We also evaluated two alternative models for combining word embeddings into sentence embeddings. Multiplicative (Mult) embeddings were constructed by adding one to each element of the word embeddings (to avoid negative numbers), then performing elementwise multiplication of all word embeddings. Convolutional (Conv) embeddings were constructed by adding one to each element of the word embeddings, then iteratively performing circular discrete convolution of each word embedding with the convolution of all previous word embeddings. For all three models based on word embeddings, sentence embeddings were constructed after removing a list of stop words containing words with little semantic content such as pronouns, modal verbs, conjunctions, and common prepositions. Cosine similarity was used to compute the similarity of each pair of sentence embeddings.

#### 3.1.3 Transformer models

We computed the representations for a range of transformer architectures, along with the older InferSent LSTM model for comparison, as summarised in Table 2. As per our preregistration, for the statistical analysis we averaged the RSA correlation with brain representations over five different transformer architectures: ERNIE 2.0, AMRBart, SentBERT, DefSent, and OpenAI. For all transformers, sentence embeddings were normalised by subtracting the mean and dividing by the standard deviation of each feature. This is motivated by research indicating that without normalisation, transformers tend to learn very anisotropic embeddings with a few dimensions dominating over all the others^80,81^. Sentence similarities were computed using cosine similarity.

**Table 2:**
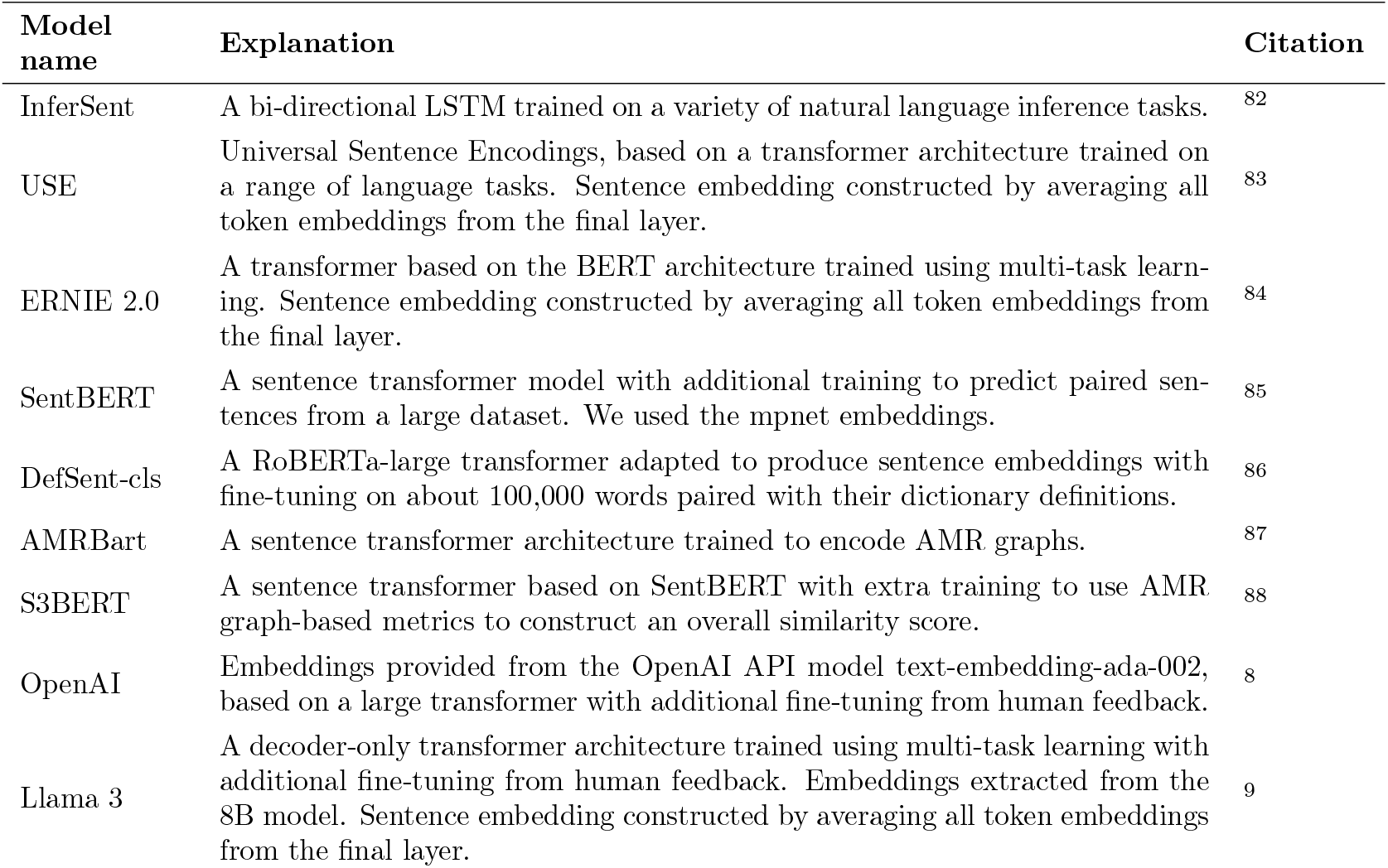
Summary of models of sentence meaning analysed in this study.

#### 3.1.4 Graph models

We adopted AMR as the primary graph-based approach for representing sentence meaning. We used the SapienzaNLP (Spring) AMR parser^89^ to parse all sentences, as it is among the best-performing AMR parsers with freely available and easily implementable code. Evaluating syntax-based models using STS datasets requires a method for computing the similarity between the graphs for each sentence. While various techniques have been developed for converting graphs into vector embeddings, these have typically focused on knowledge databanks rather than natural language^90,91^. Furthermore, we are interested in testing graph-based models of representing sentences more directly, rather than the embeddings produced from these graphs. As such, we analyse the similarity of AMR graphs using two existing methods for comparing graph similarity directly: SMATCH^92^ and WWLK^93^, yielding the AMR-Smatch and AMR-WLK models respectively. The SMATCH metric computes the number of matching triples (sets of three connected nodes) that two AMR graphs share in common relative to the total number of triples across both graphs. The WWLK metric uses a very different approach, first constructing a vector embedding for each node based on its connections to other nodes, then concatenating across all nodes in the graph, and finally computing the transformation distance between these two concatenated node vectors. In the main manuscript we report the results for the more widely-used SMATCH metric, as it achieved much higher correlations than the WWLK metric.

As a supplementary analysis, we also evaluated constituency parses produced using two different methods. In the first, we constructed a simple context-free grammar (CFG) to produce candidate parses of all sentences, with the most plausible parse manually selected from these candidates. In the second approach, we used the Berkeley Neural Parser as implemented in the benepar python library to automatically parse all sentences^94,95^. To compare the similarity of these graphs, we used both the edit distance and subtree similarity metrics^96^.

#### 3.1.5 Hybrid models

To compute representations for the VerbNet-CN hybrid model, we used the GPT-4 model of the OpenAI Chat Completions API to parse each of the 108 sentences by assigning parts of the sentence to one of eight semantic roles: Verb, Agent, Patient, Theme, Time, Manner, Location, Trajectory. After parsing by semantic role, we then computed embeddings for each semantic role as be-fore, by averaging the static ConceptNet embeddings of each constituent word after the removal of stop words. Words that are not associated with any semantic role were discarded. As before, the result is a set of role embeddings which constitutes the representation of the meaning of the sentence in terms of vector representations of each major semantic role.

To compute the similarity between two sentences, we first aligned the two sentences based on the semantic roles. Matching semantic roles were then accorded a similarity of 0.5 plus the computed cosine similarity between the rolewise embeddings. In cases where the semantic role was present in one sentence but not the other, a rolewise similarity of zero was used. Overall sentence similarity was computed as the weighted average of these eight rolewise similarities. We used fixed weights of 3 for the Verb, and 2 for Agent, Patient, and Theme, and 0.5 for Time, Manner, Location, and Trajectory, adopted from our previous study^31^.

To compute representations for the AMR-CN hybrid model^31^, we first parsed sentences using the SapienzaNLP (Spring) AMR parser^89^. Each token in the sentence was then assigned an ‘AMR role’ in accordance with its location in the parse tree by concatenating all nested parse labels. Role similarities were computed as the cosine similarity between the averaged ConceptNet word embeddings for all tokens with the same AMR role in each sentence of a sentence pair. Finally, the overall sentence similarity was computed as average role similarity over all roles found.

### 3.2 fMRI data collection

#### 3.2.1 Participants

Thirty-nine participants (23 women, 14 men, 2 other) between the ages of 18 and 40 (mean=22.2) were recruited from our university campus (The University of Melbourne) for the study. All self-identified as native speakers of English, and all but one (a last-minute replacement) identified as right-handed. Participants received $70 as compensation for their time, which corresponds to about $23 per hour for a three-hour session. Nine participants were excluded from the main analysis: seven for scoring below 70% on the attention task (see details below), and two for head motion exceeding 4mm maximum framewise deviation averaged over eight runs, leaving data from 30 participants for subsequent analysis. Note that owing to somewhat poorer performance of participants compared to those in our pilot, we lowered the cutoff slightly from the 75% stated in the preregistration, which led to the inclusion of a single additional participant who scored 73%. In Figure S19 we show that accuracy on attention check questions had a strong association with model correlations in line with our expectations.

The study protocol was approved by the University of Melbourne Human Research Ethics Committee (Reference Number: 2023-28035-47583-3).

#### 3.2.2 Experimental task

While undergoing scanning, participants were presented with a set of 108 sentences, each shown one at a time. They were instructed simply to read each sentence and think about its meaning. Sentence timing was varied with the length of the sentence, to allow sufficient time for reading longer sentences while avoiding leaving time for participants to engage in mind wandering after reading the shorter sentences. The time for each sentence was computed using a quadratic formula in the number of characters, with parameters chosen based on feedback from pilot participants. Presentation time ranged from 2-7 seconds, with an average of 4.29 seconds per sentence. The inter-stimulus interval was selected from a uniform random distribution between 2-7s, with an average of 4.5s. The order of sentences was randomised separately for each participant, with 54 sentences presented during each 508s run. The entire set of 108 sentences was presented every two runs, such that upon completion of all eight runs participants had seen each sentence four times. For five participants, only six runs were included, either because the participant did not complete the full scan or due to excessive head motion on the remaining two runs.

#### 3.2.3 Attention task

To check attention and task engagement, participants were presented with four questions randomly distributed throughout each of the eight runs (40 questions total). All questions were four-option multiple choice questions relating to the meaning of the immediately preceding sentence. Each question, along with its potential answers, was displayed on screen for 5 seconds. Participants selected the answer using one of the two-button boxes held in each hand.

#### 3.2.4 Image acquisition

The fMRI data was acquired using a 7 Tesla Siemens MAGNETOM scanner at the Melbourne Brain Centre (Parkville, Victoria) with a 32-channel radio frequency coil. The BOLD signal was measured using a multiband echo-planar imaging sequence (TR = 800 ms, TE= 22.2 ms, FA = 45°). We acquired 636 volumes on each of the eight runs, each with 84 interleaved slices (thickness = 1.6 mm, gap = 0 mm, FOV = 208mm, matrix = 130×130, multi-band factor = 6, voxel size=1.6×1.6×1.6mm^3^). Cardiac and respiratory traces were also recorded.

#### 3.2.5 Preprocessing

Preprocessing was performed using fMRIprep with default parameters^97^. First, the T1-weighted (T1w) structural image was skull-stripped and normalised to the MNI152NLin2009cAsym standard space. Second, each of the 8 BOLD runs was slice-time corrected and the volumes were motion-corrected by registering them to the single-band reference (SBRef) for each run. Distortion correction was applied by mapping field coefficients onto the reference image. All BOLD runs were then coregistered to the T1w reference, and resampled into the standard 1.6mm MNI152NLin2009cAsym space. Full details of this process are given in Supplementary Information.

#### 3.2.6 GLM Model

To model the brain activity pattern resulting from each sentence, a general linear model (GLM) was fitted using a boxcar function for each separate sentence convolved with the canonical haemodynamic response (HRF). This approach yields beta coefficients for each voxel and each distinct sentence stimulus. GLMs were fitted using GLM-single^98^, a sophisticated software package able to automatically detect and remove sources of noise, and also fit an appropriate HRF for each voxel.

A constant stimulus duration of 3s was used for all stimuli for two reasons. First, GLMsingle does not support variable stimuli lengths. Second, participants will not form a full mental representation of a sentence until they finish reading it, so it is appropriate to only include the final portion of the stimulus for longer sentences.

In our preregistration we stated we would extract the representation over the final 3s for each stimulus. However, during the course of the study it became clear from participant feedback that the time provided for reading longer sentences was more than necessary, particularly for repeated trials. As such, in the main manuscript we instead report results for the middle 3s of each stimulus. For example, for a 7s sentence representations are evaluated during the window 2-5s. We show in Figure S20 that our results are similar when using the final 3s but with lower absolute magnitudes, presumably because participants begin to disengage with the task at the end of longer sentence presentations.

Three regressors of no interest were included in the GLM. The first was the number of characters displayed to the participant at any given time, as a control for the optical size of the visual stimulus. The final two regressors specified the timing of button presses for question responses, with one regressor each for left-hand and right-hand presses.

Regressions were run for each subject using the default parameters. Beta coefficients for each presentation of all 108 sentences were then extracted from the final ‘TYPED FITHRF GLMDENOISE’ output of GLMsin-gle, and averaged over all four presentations of each sentence.

### 3.3 Behavioural data collection

#### 3.3.1 Participants

A total of 502 participants (267 male, 223 female, and 17 other; age range, 18-45; mean age ± SD, 29.80 ± 6.0) were recruited using the Prolific platform (https://www.prolific.com/). Participants were paid £4.50 for completing the task, which took an average of 22.5 minutes, amounting to an hourly rate of £11.96. All participants were self-declared native English speakers in Australia or the United States.

The study protocol was approved by the University of Melbourne Human Research Ethics Committee (Reference Number: 2023-23559-36378-6).

#### 3.3.2 Survey task

Each participant provided similarity judgements on a 7-point Likert scale (1-7) of 102 sentence pairs randomly selected from the pool of all 5,778 sentence pairs. As our primary interest was in the block diagonal sentences, we over-sampled from these sentence pairs relative to the other sentence pairs. As such, each participant rated 42 block diagonal sentence pairs and 60 other sentence pairs. Given the inherent vagueness of the similarity judgement task, previous studies have noted that lengthy instructions on how to make similarity judgements are often unclear, or may bias participant responses^99,100^. Because our goal was to elicit intuitive judgements without imposing any particular framework which might influence results, we did not provide participants with any special training or instructions about how to assign ratings. Participants were simply instructed to “consider both the similarity in meaning of the individual words contained in the sentences, as well as the similarity of the overall idea or meaning expressed by the sentences”. The full instructions given to participants can be found in the Supplementary Information.

In addition to the sentence pairs derived from the 108 experimental sentences, participants were also presented with additional 10 sentence pairs that served as an attention check. These stimuli consisted of either pairs of identical sentences (high similarity) or one simple sentence paired with a grammatically correct but nonsensical sentence (low similarity).

#### 3.3.3 Preprocessing

We excluded all participants who failed more than two of the ten attention check items, resulting in 486 of 502 participants being retained. This amounted to 49,572 judgements, providing an average of 22 ratings for each block diagonal sentence pair and 6 for each of the other sentence pairs. Similarity judgements were averaged over participants and normalised between 0 and 1.

#### 3.3.4 GPT-4 ratings

As an additional comparison to human similarity judgements, we also collected similarity ratings using the API of the GPT-4 model^101^. We passed each distinct pair of 5,778 sentences to GPT-4 one at a time, to avoid any spurious effects of recent context. The prompt we used is given below:

“You will be presented with two sentences. Your task is to judge how similar is the meaning of the two sentences. You will make this judgement by choosing a rating from 0 (most dissimilar) to 1 (most similar) to two decimal places. In providing your rating, consider both the similarity in meaning of the individual words contained in the sentences, as well as the similarity of the overall idea or meaning expressed by the sentences. Provide a numerical rating only; do not explain your answers. Here are the sentences:”

### 3.4 Representational Similarity Analysis

#### 3.4.1 Voxel Selection

Voxel selection was performed in two different ways. To provide an overall brain representation, we extracted all voxels within the cortical mask from the MNI152NLin2009cAsym template. To eliminate potential confound from visual regions, we also constructed a cortical mask excluding voxels in visual cortical regions V1-V4 from the cortical mask. In our preregistration we stated that we would remove any voxels having an absolute correlation with sentence length greater than 0.5. However during our analysis we found this to be infeasible given the large number of voxels sensitive to sentence length. We subsequently became aware that several previous studies have found similar length effects in the language network^71–73^. As such, we instead directly remove the visual cortex regions V1-V4 from analysis. As an additional check, we also performed all analyses controlling for the minimum sentence length, with the results shown in Figure S4. In addition, we also analysed voxels within a language region of interest (ROI) mask. The mask contains 26,000 voxels found to be primarily sensitive to linguistic stimuli in a series of previous experiments involving contrasting sentence stimuli with pseudowords^15^.

To identify voxels sensitive to sentence stimuli, the stability score was computed for each voxel as the average correlation between its time series of activity on different presentations of the stimuli^2^. All voxels within the mask with stability scores above a threshold of 0.07 were selected for computing RSA matrices. We show in Figure S18 that alternative stability thresholds yield similar results, though with higher magnitudes when higher thresholds are used.

Masks for cortical regions of interest were constructed using the Glasser atlas^102^. Parcel indices included in each region were as follows. Dorsolateral prefrontal cortex: 67,68,71,73,83,84,85,86,87; dorsomedial prefrontal cortex: 26,43,63,69; precuneus: 15,27,29,30,31,45,121,142; posterior cingulate: 14,32,33,34,35,38,161,162; primary visual cortex: 1,4,5,6; primary somatosensory cortex: 9,51,52,53.

#### 3.4.2 Computing RSA matrices

For fMRI data, RSA matrices were computed by first normalising GLMsingle beta coefficients by subtracting the mean and dividing by the standard deviation for each voxel. Cosine similarities were then computed between the voxel representations of each sentence (using only the subset of included voxels) for each distinct pair of sentences, yielding an RSA matrix for each participant.

RSA matrices for computational models were computed differently depending on the model in question. For all vector-based models (including Mean and Trans-former) sentence embeddings were extracted for each sentence, and then normalised by subtracting the mean and dividing by the standard deviation for each dimension. Pairwise sentence similarities were then computed using cosine similarity between the corresponding embeddings. As an additional check, for the vector-based models we also computed similarities using the Dimension Insensitive Euclidean Metric (DIEM), which is designed to adjust for the effects of differences in dimensionality be-tween embedding models^103^.

For models not entirely based on vector representations (i.e. the constituency parsers, the AMR-based models, and VerbNet-ConceptNet), we compute pairwise similarities as specified in subsection 3.1.

#### 3.4.3 Data-model RSA correlations

RSA matrices for brain representations were compared with those of the computational models by calculating for each participant the partial Spearman correlation controlling for the difference in sentence lengths, then averaging over participants. We use the pingouin 0.5.4 python package, which utilises the inverse covariance matrix for computing partial Spearman correlations. This has been proven more reliable than the alternative regression residuals technique when a subset of variables are discrete (see discussion at https://github.com/raphaelvallat/pingouin/issues/147). This is especially relevant for the AMR-Smatch model, as the Smatch metric outputs a discrete similarity score.

In addition to the simple average across participants, we implemented an alternative method adapted from several previous studies^26,27^, in which a group-averaged RSA matrix was first constructed by averaging pairwise sentence similarities over participants, and then the correlation computed between each model RSA and this group-averaged RSA matrix.

For the simple average method, confidence intervals and statistical testing was performed using simple two-sided t-tests computed over participants. For the group average method, confidence intervals were estimated by bootstrapping over participants, performed 100 times. In the preregistration we planned to perform bootstrapping over stimuli as well as over participants, however in retrospect we judged this to be inappropriate since our sentences were not a random sampling from some corpus, but were specially constructed to provide specified semantic and syntactic variation. For both methods, the Bonferroni correction was used to adjust for three independent model comparisons (Mean to Transformer, Transformer to VerbNet-CN, and VerbNet-CN to AMR-Smatch), yielding a significance level of *α*=0.05/3 = 0.0167.

We also computed the correlation between humanrated similarities and the brain RSA similarities, though we did not perform a statistical test as we had no prior hypothesis about this correlation.

### 3.5 Searchlight RSA

To visualise the location of the cortical regions responsible for encoding sentence information, we implemented RSA-searchlight^104^. Using the mne-rsa pack-age (see https://users.aalto.fi/~vanvlm1/mne-rsa/index.html), we performed an 8mm searchlight analysis over all voxels within the cortical mask. Images were smoothed with 5mm FWHM and thresholded at z=2 using threshold-free cluster enhancement (TFCE) correction for display.

## Notes

https://openneuro.org/datasets/ds007393

https://github.com/Fods12/sentence_meaning_in_the_brain

### Competing Interest Statement

The authors have declared no competing interest.

### Summary of Updates

New SI figures, modified conclusion, fixed typographical errros.

## References

1. Mitchell, T. et al. Predicting human brain activity associated with the meanings of nouns. Science 320, 1191–1195. ISSN: 0036-8075 (2008).

2. Just, M., Cherkassky, V. L., Aryal, S. & Mitchell, T. M. A neurosemantic theory of concrete noun representation based on the underlying brain codes. PloS one 5, e8622. ISSN: 1932-6203 (2010).

3. Wehbe, L. et al. Simultaneously uncovering the patterns of brain regions involved in different story reading subprocesses. PloS one 9, e112575. ISSN: 1932-6203 (2014).

4. Huth, A. G., De Heer, W. A., Griffiths, T. L., Theunissen, F. E. & Gallant, J. L. Natural speech reveals the semantic maps that tile human cerebral cortex. Nature 532, 453–458. ISSN: 1476-4687 (2016).

5. Pereira, F. et al. Toward a universal decoder of linguistic meaning from brain activation. Nature communications 9, 1–13. ISSN: 2041-1723 (2018).

6. Günther, F., Rinaldi, L. & Marelli, M. Vector-space models of semantic representation from a cognitive perspective: A discussion of common mis-conceptions. Perspectives on Psychological Science 14, 1006–1033. ISSN: 1745-6916 (2019).

7. Karamolegkou, A., Abdou, M. & Søgaard, A. Mapping Brains with Language Models: A Survey. arXiv preprint arXiv:2306.05126 (2023).

8. Ouyang, L. et al. Training language models to follow instructions with human feedback. Advances in Neural Information Processing Systems 35, 27730–27744 (2022).

9. Touvron, H. et al. Llama: Open and efficient foundation language models. arXiv preprint arXiv:2302.13971 (2023).

10. Team, G. et al. Gemini: a family of highly capable multimodal models. arXiv preprint arXiv:2312.11805 (2023).

11. Bhatia, S., Richie, R. & Zou, W. Distributed semantic representations for modeling human judgment. Current Opinion in Behavioral Sciences 29, 31–36 (2019).

12. Erk, K. The probabilistic turn in semantics and pragmatics. Annual Review of Linguistics 8, 101–121 (2022).

13. Tuckute, G., Kanwisher, N. & Fedorenko, E. Language in brains, minds, and machines. Annual Review of Neuroscience 47, 277–301 (2024).

14. Anderson, A. et al. Deep artificial neural networks reveal a distributed cortical network encoding propositional sentence-level meaning. Journal of Neuroscience 41, 4100–4119. ISSN: 0270-6474 (2021).

15. Schrimpf, M. et al. The neural architecture of language: Integrative modeling converges on predictive processing. Proceedings of the National Academy of Sciences 118. ISSN: 0027-8424 (2021).

16. Antonello, R., Turek, J. S., Vo, V. & Huth, A. Low-dimensional structure in the space of language representations is reflected in brain responses. Advances in neural information processing systems 34, 8332–8344 (2021).

17. Pasquiou, A., Lakretz, Y., Hale, J. T., Thirion, B. & Pallier, C. Neural Language Models are not Born Equal to Fit Brain Data, but Training Helps in International Conference on Machine Learning (PMLR, 2022), 17499–17516.

18. Aliko, S., Huang, J., Gheorghiu, F., Meliss, S. & Skipper, J. I. A naturalistic neuroimaging database for understanding the brain using ecological stimuli. Scientific Data 7, 1–21. ISSN: 2052-4463 (2020).

19. Hamilton, L. S. & Huth, A. G. The revolution will not be controlled: natural stimuli in speech neuroscience. Language, cognition and neuroscience 35, 573–582. ISSN: 2327-3798 (2020).

20. Nastase, S. A. et al. The “Narratives” fMRI dataset for evaluating models of naturalistic language comprehension. Scientific data 8, 250 (2021).

21. Zhang, Y., Kim, J.-H., Brang, D. & Liu, Z. Naturalistic stimuli: A paradigm for multiscale functional characterization of the human brain. Current opinion in biomedical engineering 19, 100298. ISSN: 2468-4511 (2021).

22. Li, J., Lai, M. & Pylkkänen, L. Semantic composition in experimental and naturalistic paradigms. Imaging Neuroscience 2, 1–17 (2024).

23. Kriegeskorte, N. & Douglas, P. K. Interpreting encoding and decoding models. Current opinion in neurobiology 55, 167–179 (2019).

24. Bruffaerts, R. et al. Redefining the resolution of semantic knowledge in the brain: advances made by the introduction of models of semantics in neuroimaging. Neuroscience & Biobehavioral Reviews 103, 3–13. ISSN: 0149-7634 (2019).

25. Hagoort, P. The meaning-making mechanism(s) behind the eyes and between the ears. Philosophical Transactions of the Royal Society B 375, 20190301. ISSN: 0962-8436 (2020).

26. Wang, X. et al. Organizational principles of abstract words in the human brain. Cerebral Cortex 28, 4305–4318. ISSN: 1047-3211 (2018).

27. Fernandino, L., Tong, J.-Q., Conant, L. L., Humphries, C. J. & Binder, J. R. Decoding the information structure underlying the neural representation of concepts. Proceedings of the National Academy of Sciences 119, e2108091119. ISSN: 0027-8424 (2022).

28. Tong, J. et al. A distributed network for multimodal experiential representation of concepts. Journal of Neuroscience 42, 7121–7130 (2022).

29. Acunzo, D. J., Low, D. M. & Fairhall, S. L. Deep neural networks reveal topic-level representations of sentences in medial prefrontal cortex, lateral anterior temporal lobe, precuneus, and angular gyrus. NeuroImage 251, 119005. ISSN: 1053-8119 (2022).

30. Fairhall, S. L. Sentence-level embeddings reveal dissociable word-and sentence-level cortical representation across coarse-and fine-grained levels of meaning. Brain and Language 250, 105389 (2024).

31. Fodor, J., De Deyne, S. & Suzuki, S. Compositionality and Sentence Meaning: Comparing Semantic Parsing and Transformers on a Challenging Sentence Similarity Dataset. Computational Linguistics, 1–52 (2024).

32. Lake, B. M. & Murphy, G. L. Word meaning in minds and machines. Psychological Review 130, 401–31. ISSN: 1939-1471 (2021).

33. Žabokrtský, Z., Zeman, D. & Ševčíková, M. Sentence meaning representations across languages: what can we learn from existing frameworks? Computational Linguistics 46, 605–665 (2020).

34. Anderson, A., Kiela, D., Clark, S. & Poesio, M. Visually grounded and textual semantic models differentially decode brain activity associated with concrete and abstract nouns. Transactions of the Association for Computational Linguistics 5, 17–30 (2017).

35. Just, M., Wang, J. & Cherkassky, V. L. Neural representations of the concepts in simple sentences: Concept activation prediction and context effects. Neuroimage 157, 511–520. ISSN: 1053-8119 (2017).

36. Fedorenko, E., Ivanova, A. A. & Regev, T. I. The language network as a natural kind within the broader landscape of the human brain. Nature Reviews Neuroscience 25, 289–324 (2024).

37. Handjaras, G. et al. How concepts are encoded in the human brain: a modality independent, category-based cortical organization of semantic knowledge. Neuroimage 135, 232–242. ISSN: 1053-8119 (2016).

38. Shain, C. et al. Distributed sensitivity to syntax and semantics throughout the language network. Journal of Cognitive Neuroscience 36, 1427–1471 (2024).

39. Fedorenko, E., Blank, I. A., Siegelman, M. & Mineroff, Z. Lack of selectivity for syntax relative to word meanings throughout the language network. Cognition 203, 104348 (2020).

40. Marelli, M. et al. A SICK cure for the evaluation of compositional distributional semantic models in Proceedings of the Ninth International Conference on Language Resources and Evaluation (LREC’14) (2014), 216–223.

41. Goucha, T. & Friederici, A. D. The language skeleton after dissecting meaning: A functional segregation within Broca’s area. NeuroImage 114, 294–302 (2015).

42. Mollica, F. et al. Composition is the core driver of the language-selective network. Neurobiology of Language 1, 104–134. ISSN: 2641-4368 (2020).

43. Fedorenko, E., Nieto-Castanon, A. & Kanwisher, N. Lexical and syntactic representations in the brain: an fMRI investigation with multi-voxel pattern analyses. Neuropsychologia 50, 499–513 (2012).

44. Matchin, W., Hammerly, C. & Lau, E. The role of the IFG and pSTS in syntactic prediction: Evidence from a parametric study of hierarchical structure in fMRI. cortex 88, 106–123 (2017).

45. Călinescu, L., Ramchand, G. & Baggio, G. How (not) to look for meaning composition in the brain: A reassessment of current experimental paradigms. Frontiers in Language Sciences 2, 1096110 (2023).

46. Frankland, S. M. & Greene, J. D. An architecture for encoding sentence meaning in left mid-superior temporal cortex. Proceedings of the National Academy of Sciences 112, 11732–11737. ISSN: 0027-8424 (2015).

47. Wang, J., Cherkassky, V., Yang, Y., Diana, N. & Just, M. A. Identifying thematic roles from neural representations measured by functional magnetic resonance imaging. Cognitive neuropsychology 33, 257–264. ISSN: 0264-3294 (2016).

48. Frankland, S. M. & Greene, J. D. Two ways to build a thought: distinct forms of compositional semantic representation across brain regions. Cerebral Cortex 30, 3838–3855. ISSN: 1047-3211 (2020).

49. Giglio, L., Hagoort, P. & Ostarek, M. Neural encoding of semantic structures during sentence production. Cerebral Cortex 34 (2024).

50. Graves, W. W., Binder, J. R., Desai, R. H., Conant, L. L. & Seidenberg, M. S. Neural correlates of implicit and explicit combinatorial semantic processing. Neuroimage 53, 638–646. ISSN: 1053-8119 (2010).

51. Schell, M., Zaccarella, E. & Friederici, A. D. Differential cortical contribution of syntax and semantics: An fMRI study on two-word phrasal processing. Cortex 96, 105–120 (2017).

52. Fyshe, A., Sudre, G., Wehbe, L., Rafidi, N. & Mitchell, T. M. The lexical semantics of adjective– noun phrases in the human brain. Human brain mapping 40, 4457–4469 (2019).

53. Ciapparelli, M., Marelli, M., Graves, W. & Reverberi, C. Compositionality in the semantic network: a model-driven representational similarity analysis. Cerebral Cortex 35, bhaf246 (2025).

54. Oota, S. R., Gupta, M. & Toneva, M. Joint processing of linguistic properties in brains and language models in Advances in Neural Information Processing Systems 36 (2024).

55. Feghhi, E., Hadidi, N., Song, B., Blank, I. A. & Kao, J. C. What Are Large Language Models Mapping to in the Brain? a Case Against Over-Reliance on Brain Scores. arXiv preprint arXiv:2406.01538 (2024).

56. Hosseini, E. A. et al. Artificial Neural Network Language Models Predict Human Brain Responses to Language Even After a Developmentally Realistic Amount of Training. Neurobiology of Language 5, 43–63. ISSN: 2641-4368. 10.1162/nol%5C_a%5C_00137 (Apr. 2024).

57. Caucheteux, C., Gramfort, A. & King, J.-R. Disentangling syntax and semantics in the brain with deep networks in International conference on machine learning (2021), 1336–1348.

58. Kauf, C., Tuckute, G., Levy, R., Andreas, J. & Fedorenko, E. Lexical-semantic content, not syntactic structure, is the main contributor to ANN-brain similarity of fMRI responses in the language network. Neurobiology of Language 5, 7–42. ISSN: 2641-4368 (2024).

59. Hadidi, N., Feghhi, E., Song, B. H., Blank, I. A. & Kao, J. C. Illusions of Alignment Between Large Language Models and Brains Emerge From Fragile Methods and Overlooked Confounds. bioRxiv, 2025–03 (2025).

60. Chang, T. A. & Bergen, B. K. Language model behavior: A comprehensive survey. Computational Linguistics 50, 293–350 (2024).

61. Clark, K., Khandelwal, U., Levy, O. & Manning, C. D. What Does BERT Look at? an Analysis of BERT’s Attention in Proceedings of the 2019 ACL Workshop BlackboxNLP: Analyzing and Interpreting Neural Networks for NLP (2019), 276–286.

62. Manning, C. D., Clark, K., Hewitt, J., Khandelwal, U. & Levy, O. Emergent linguistic structure in artificial neural networks trained by self-supervision. Proceedings of the National Academy of Sciences 117, 30046–30054 (2020).

63. Opitz, J. & Frank, A. Better Smatch= Better Parser? AMR evaluation is not so simple anymore in Proceedings of the 3rd Workshop on Evaluation and Comparison of NLP Systems (2022), 32–43.

64. Leung, W. C., Wein, S. & Schneider, N. Semantic Similarity as a Window into Vector-and Graph-Based Metrics in Proceedings of the 2nd Workshop on Natural Language Generation, Evaluation, and Metrics (GEM) (2022), 106–115.

65. Opitz, J. SMATCH++: Standardized and Extended Evaluation of Semantic Graphs in Findings of the Association for Computational Linguistics: EACL 2023 (2023), 1595–1607.

66. Hamilton, J. P. et al. Default-mode and task-positive network activity in major depressive disorder: implications for adaptive and maladaptive rumination. Biological psychiatry 70, 327–333 (2011).

67. Kuhnke, P., Kiefer, M. & Hartwigsen, G. Conceptual representations in the default, control and attention networks are task-dependent and cross-modal. Brain and Language 244, 105313 (2023).

68. Fernandino, L. & Binder, J. R. How does the “default mode” network contribute to semantic cognition? Brain and Language 252, 105405 (2024).

69. Blank, I. A. & Fedorenko, E. No evidence for differences among language regions in their temporal receptive windows. NeuroImage 219, 116925 (2020).

70. Brennan, J. R., Stabler, E. P., Van Wagenen, S. E., Luh, W.-M. & Hale, J. T. Abstract linguistic structure correlates with temporal activity during naturalistic comprehension. Brain and language 157, 81–94. ISSN: 0093-934X (2016).

71. Nelson, M. J. et al. Neurophysiological dynamics of phrase-structure building during sentence processing. Proceedings of the National Academy of Sciences 114, E3669–E3678. ISSN: 0027-8424 (2017).

72. Schuster, S., Hawelka, S., Himmelstoss, N. A., Richlan, F. & Hutzler, F. The neural correlates of word position and lexical predictability during sentence reading: Evidence from fixation-related fMRI. Language, Cognition and Neuroscience 35, 613–624 (2020).

73. Woolnough, O. et al. Spatiotemporally distributed frontotemporal networks for sentence reading. Proceedings of the National Academy of Sciences 120, e2300252120. ISSN: 0027-8424 (2023).

74. Deniz, F., Tseng, C., Wehbe, L., la Tour, T. D. & Gallant, J. L. Semantic representations during language comprehension are affected by context. Journal of Neuroscience 43, 3144–3158. ISSN: 0270-6474 (2023).

75. Baron, S. G., Thompson-Schill, S. L., Weber, M. & Osherson, D. An early stage of conceptual combination: Superimposition of constituent concepts in left anterolateral temporal lobe. Cognitive neuroscience 1, 44–51 (2010).

76. Frankland, S. M. & Greene, J. D. Concepts and compositionality: in search of the brain’s language of thought. Annual review of psychology 71, 273–303. ISSN: 0066-4308 (2020).

77. Desbordes, T., King, J.-R. & Dehaene, S. Tracking the neural codes for words and phrases during semantic composition, working-memory storage, and retrieval. Cell Reports 43 (2024).

78. Stanojević, M., Brennan, J. R., Dunagan, D., Steedman, M. & Hale, J. T. Modeling structure-building in the brain with CCG parsing and large language models. Cognitive science 47, e13312 (2023).

79. Fresen, A. J., Choenni, R., Heilbron, M., Zuidema, W. & de Heer Kloots, M. Language Models That Accurately Represent Syntactic Structure Exhibit Higher Representational Similarity To Brain Activity in Proceedings of the Annual Meeting of the Cognitive Science Society 46 (2024).

80. Timkey, W. & van Schijndel, M. All Bark and No Bite: Rogue Dimensions in Transformer Language Models Obscure Representational Quality in Proceedings of the 2021 Conference on Empirical Methods in Natural Language Processing (2021), 4527–4546.

81. Cai, X., Huang, J., Bian, Y. & Church, K. Isotropy in the contextual embedding space: Clusters and manifolds in International Conference on Learning Representations (2021).

82. Conneau, A., Kiela, D., Schwenk, H., Barrault, L. & Bordes, A. Supervised Learning of Universal Sentence Representations from Natural Language Inference Data in Proceedings of the 2017 Conference on Empirical Methods in Natural Language Processing (Association for Computational Linguistics, Copenhagen, Denmark, Sept. 2017), 670–680. https://aclanthology.org/D17-1070.

83. Cer, D. et al. Universal sentence encoder for English in Proceedings of the 2018 conference on empirical methods in natural language processing: system demonstrations (2018), 169–174.

84. Sun, Y. et al. Ernie 2.0: A continual pre-training framework for language understanding. Proceedings of the AAAI conference on artificial intelligence 34, 8968–8975 (2020).

85. Reimers, N. & Gurevych, I. Sentence-BERT: Sentence Embeddings using Siamese BERT-Networks in Proceedings of the 2019 Conference on Empirical Methods in Natural Language Processing and the 9th International Joint Conference on Natural Language Processing (EMNLP-IJCNLP) (2019), 3982–3992.

86. Tsukagoshi, H., Sasano, R. & Takeda, K. Def-Sent: Sentence Embeddings using Definition Sentences in Proceedings of the 59th Annual Meeting of the Association for Computational Linguistics and the 11th International Joint Conference on Natural Language Processing (Volume 2: Short Papers) (2021), 411–418.

87. Bai, X., Chen, Y. & Zhang, Y. Graph Pre-training for AMR Parsing and Generation in Proceedings of the 60th Annual Meeting of the Association for Computational Linguistics (Volume 1: Long Papers) (2022), 6001–6015.

88. Opitz, J. & Frank, A. SBERT studies Meaning Representations: Decomposing Sentence Embeddings into Explainable Semantic Features in Proceedings of the 2nd Conference of the Asia-Pacific Chapter of the Association for Computational Linguistics and the 12th International Joint Conference on Natural Language Processing (Volume 1: Long Papers) (2022), 625–638.

89. Bevilacqua, M., Blloshmi, R. & Navigli, R. One SPRING to Rule Them Both: Symmetric AMR Semantic Parsing and Generation without a Complex Pipeline in Proceedings of AAAI (2021), 12564–12573.

90. Goyal, P. & Ferrara, E. Graph embedding techniques, applications, and performance: A survey. Knowledge-Based Systems 151, 78–94 (2018).

91. Rossi, A., Barbosa, D., Firmani, D. Matinata, & Merialdo, P. Knowledge graph embedding for link prediction: A comparative analysis. ACM Transactions on Knowledge Discovery from Data (TKDD) 15, 1–49 (2021).

92. Cai, S. & Knight, K. Smatch: an evaluation metric for semantic feature structures in Proceedings of the 51st Annual Meeting of the Association for Computational Linguistics (Volume 2: Short Papers) (2013), 748–752.

93. Opitz, J., Daza, A. & Frank, A. Weisfeiler-leman in the bamboo: Novel AMR graph metrics and a benchmark for AMR graph similarity. Transactions of the Association for Computational Linguistics 9, 1425–1441 (2021).

94. Kitaev, N. & Klein, D. Constituency Parsing with a Self-Attentive Encoder in Proceedings of the 56th Annual Meeting of the Association for Computational Linguistics (Volume 1: Long Papers) (Association for Computational Linguistics, Melbourne, Australia, July 2018), 2676–2686. https://www.aclweb.org/anthology/P18-1249.

95. Kitaev, N. & Klein, D. Berkeley Neural Parser 2021. https://github.com/nikitakit/self-attentive-parser.

96. Collins, M. & Duffy, N. Convolution kernels for natural language. Advances in neural information processing systems 14 (2001).

97. Esteban, O. et al. fMRIPrep: a robust preprocessing pipeline for functional MRI. Nature Methods 16, 111–116 (2019).

98. Prince, J. S. et al. Improving the accuracy of single-trial fMRI response estimates using GLM-single. Elife 11, e77599. ISSN: 2050-084X (2022).

99. Abe, K., Yokoi, S., Kajiwara, T. & Inui, K. Why is sentence similarity benchmark not predictive of application-oriented task performance? in Proceedings of the 3rd Workshop on Evaluation and Comparison of NLP Systems (2022), 70–87.

100. Abdalla, M., Vishnubhotla, K. & Mohammad, S. What Makes Sentences Semantically Related? A Textual Relatedness Dataset and Empirical Study in Proceedings of the 17th Conference of the European Chapter of the Association for Computational Linguistics (2023), 782–796.

101. Achiam, J. et al. GPT-4 Technical Report. arXiv preprint arXiv:2303.08774 (2023).

102. Glasser, M. F. et al. A multi-modal parcellation of human cerebral cortex. Nature 536, 171–178 (2016).

103. Tessari, F., Yao, K. & Hogan, N. Surpassing Cosine Similarity for Multidimensional Comparisons: Dimension Insensitive Euclidean Metric. arXiv preprint (2024).

104. Nili, H. et al. A toolbox for representational similarity analysis. PLoS computational biology 10, e1003553 (2014).

