## Supplementary Information for "When word order matters: human brains represent sentence meaning differently from large language models"

### 1 Supplementary Information

#### 1.1 Supplementary methods

##### 1.1.1 Full fMRIprep methods

Results included in this manuscript come from preprocessing performed using *fMRIPrep* 23.2.3[1,2], which is based on *Nipype* 1.8.6[3,4]. Many internal operations of *fMRIPrep* use *Nilearn* 0.10.2[5], mostly within the functional processing workflow. For more details of the pipeline, see the section corresponding to workflows in *fMRIPrep*'s documentation.

**Preprocessing of B0 inhomogeneity mappings** A total of 8 fieldmaps were found available within the input BIDS structure for this particular subject. A *B0*-nonuniformity map (or *fieldmap*) was estimated based on two (or more) echo-planar imaging (EPI) references with `topup`[6].

**Anatomical data preprocessing** A total of 1 T1-weighted (T1w) images were found within the input BIDS dataset. The T1w image was corrected for intensity non-uniformity (INU) with `N4BiasFieldCorrection`[7], distributed with ANTs 2.5.0[8], and used as T1w-reference throughout the workflow. The T1w-reference was then skull-stripped with a *Nipype* implementation of the `antsBrainExtraction.sh` workflow (from ANTs), using OASIS30ANTs as target template. Brain tissue segmentation of cerebrospinal fluid (CSF), white-matter (WM) and gray-matter (GM) was performed on the brain-extracted T1w using `fast` FSL fsl˙fast. Brain surfaces were reconstructed using `recon-all` FreeSurfer 7.3.2[9], and the brain mask estimated previously was refined with a custom variation of the method to reconcile ANTs-derived and FreeSurfer-derived segmentations of the cortical gray-matter of Mindboggle[10]. Volume-based spatial normalization to one standard space (MNI152NLin2009cAsym) was performed through nonlinear registration with `antsRegistration` ANTs 2.5.0, using brain-extracted versions of both T1w reference and the T1w template. The following template was were selected for spatial normalization and accessed with *TemplateFlow*[11]: *ICBM 152 Nonlinear Asymmetrical template version 2009c*[12], TemplateFlow ID: MNI152NLin2009cAsym.

**Functional data preprocessing** For each of the 8 BOLD runs found per subject (across all tasks and sessions), the following preprocessing was performed. First, a reference volume was generated, using a custom methodology of *fMRIPrep*, for use in head motion correction. Head-motion parameters with respect to the BOLD reference (transformation matrices, and six corresponding rotation and translation parameters) are estimated before any spatiotemporal filtering using `mcflirt` FSL[13]. The estimated *fieldmap* was then aligned with rigid-registration to the target EPI (echo-planar imaging) reference run. The field coefficients were mapped on to the reference EPI using the transform. The BOLD reference was then co-registered to the T1w reference using `bbregister` (FreeSurfer) which implements boundary-based registration[14]. Co-registration was configured with six degrees of freedom. Several confounding time-series were calculated based on the *preprocessed BOLD*: framewise displacement (FD), DVARS and three region-wise global signals. FD was computed using two formulations following Power (absolute sum of relative motions[15]) and Jenkinson (relative root mean square displacement between affines[13]). FD and DVARS are calculated for each functional run, both using their implementations in *Nipype* following published definitions[15]. The three global signals are extracted within the CSF, the WM, and the whole-brain masks. Additionally, a set of physiological regressors were extracted to allow for component-based noise correction *CompCor*[16].

Principal components are estimated after high-pass filtering the *preprocessed BOLD* time-series (using a discrete cosine filter with 128s cut-off) for the two *CompCor* variants: temporal (tCompCor) and anatomical (aCompCor). tCompCor components are then calculated from the top 2% variable voxels within the brain mask. For aCompCor, three probabilistic masks (CSF, WM and combined CSF+WM) are generated in anatomical space. The implementation differs from that of Behzadi et al. in that instead of eroding the masks by 2 pixels on BOLD space, a mask of pixels that likely contain a volume fraction of GM is subtracted from the aCompCor masks. This mask is obtained by dilating a GM mask extracted from the FreeSurfer's *aseg* segmentation, and it ensures components are not extracted from voxels containing a minimal fraction of GM. Finally, these masks are resampled into BOLD space and binarized by thresholding at 0.99 (as in the original implementation). Components are also calculated separately within the WM and CSF masks. For each CompCor decomposition, the $k$ components with the largest singular values are retained, such that the retained components' time series are sufficient to explain 50 percent of variance across the nuisance mask (CSF, WM, combined, or temporal). The remaining components are dropped from consideration.

The head-motion estimates calculated in the correction step were also placed within the corresponding confounds file. The confound time series derived from head motion estimates and global signals were expanded with the inclusion of temporal derivatives and quadratic terms for each[17]. Frames that exceeded a threshold of 0.5 mm FD or 1.5 standardized DVARS were annotated as motion outliers. Additional nuisance timeseries are calculated by means of principal components analysis of the signal found within a thin band (*crown*) of voxels around the edge of the brain, as proposed previously[18]. All resamplings can be performed with *a single interpolation step* by composing all the pertinent transformations (i.e. head-motion transform matrices, susceptibility distortion correction when available, and co-registrations to anatomical and output spaces). Gridded (volumetric) resamplings were performed using `nitransforms`, configured with cubic B-spline interpolation.

###### 1.1.2 Instructions for behavioural task

The following instructions were presented to participants in the online Prolific experiment:

> Please read the following instructions carefully before proceeding.
>
> In this questionnaire you will be presented with 112 paired sentences, in 16 groups of 7 questions per page. Your task is to judge how similar is the meaning of the two sentences. You will make this judgement by choosing a rating from 1 (very dissimilar) to 7 (very similar). In providing your rating, consider both the similarity in meaning of the individual words contained in the sentences, as well as the similarity of the overall idea or meaning expressed by the sentences.
>
> Some of the sentences may be slightly unusual or ambiguous; nevertheless you should do your best to understand their likely meaning. Bear in mind that we are not looking for any one specific 'right answer' or strategy in your responses. Your task is simply to make a judgement about how similar you think is the meaning of the two paired sentences. The only exception is that if you find a sentence that truly does not make any sense at all, then you should give it a very low similarity to whatever it is paired with. In all other cases, make your best judgement based on your assessment of overall meaning of the sentences.
>
> There is no time limit to this task, however each sentence pair should not take more than a few seconds to judge. There is no need to spend a long time pondering each sentence. In total the task should take around 20-30 minutes.
>
> Thanks very much for your time!

##### 1.2 Supplementary tables

###### 1.2.1 Complete set of stimulus sentences

| Set | Type | Sentence |
|---|---|---|
| 1 | Base | The cameraman brought the equipment to the director. |
| 1 | Same | The cameraman brought the new equipment to the director. |
| 1 | Substituted | A painter brought the equipment to the director. |
| 1 | Substituted | The cameraman sold the equipment to the director. |
| 1 | Substituted | The cameraman brought a hot lunch for the director. |
| 1 | Modified | Every morning the cameraman brought the equipment to the impatient director. |
| 1 | Modified | A team of cameramen brought all the necessary equipment to the busy director. |
| 1 | Modified | The cameraman begrudgingly brought the awkward equipment up from the basement to the director. |
| 1 | Modified | At dawn the industrious cameraman brought the essential equipment to the director in his downtown office. |
| 1 | Swapped | The director brought the cameraman to the equipment. |
| 1 | Swapped | The director brought the cameraman to his equipment. |
| 1 | Swapped | The director brought his cameraman a hot lunch. |
| 1 | Swapped | The director sold the cameraman's equipment. |
| 1 | Swapped | The director brought the painter to his equipment. |
| 1 | Swapped | Every morning the director brought the cameraman to the expensive equipment. |
| 1 | Swapped | The director inexplicably brought the cameraman to the useless old equipment in the storeroom. |
| 1 | Swapped | The renowned director brought the impatient cameraman to the equipment from his last project. |

| | | |
|---|---|---|
| 1 | Swapped | On the weekend the helpful film director brought the inexperienced cameraman to his equipment in the storeroom. |
| 2 | Base | The psychologist spoke to the secretary about the patient. |
| 2 | Same | The psychologist often spoke to the secretary about the patient. |
| 2 | Substituted | The cleaner spoke to the secretary about the patient. |
| 2 | Substituted | The psychologist warned the secretary about the patient. |
| 2 | Substituted | A psychologist spoke to her gardener about the patient. |
| 2 | Modified | The psychologist sometimes spoke to her secretary about the problematic patient. |
| 2 | Modified | The distinguished psychologist spoke in her office to the diligent secretary about her patients. |
| 2 | Modified | Occasionally the psychologist accidently spoke to his secretary about one patient within earshot of other patients. |
| 2 | Modified | One per week the psychologist spoke delicately to her experienced secretary about her most difficult patients. |
| 2 | Swapped | The secretary spoke to the patient about the psychologist. |
| 2 | Swapped | The secretary often spoke to the patient about the psychologist. |
| 2 | Swapped | The gardener spoke to the patient about her psychologist. |
| 2 | Swapped | The secretary recommended the patient to the psychologist. |
| 2 | Swapped | The secretary spoke to the patient about the cleaner. |
| 2 | Swapped | The secretary sometimes spoke rudely to patients about the psychologist. |
| 2 | Swapped | The secretary at the desk spoke clearly with the disabled patient about their next psychologist appointment. |
| 2 | Swapped | The exhausted secretary spoke slowly to the patient about their psychologist late in the evening. |
| 2 | Swapped | The concerned secretary spoke gently to the anxious patient in a private consultation room about the psychologist. |
| 3 | Base | The company purchased the apartment block from the family. |
| 3 | Same | The company bought the apartment block from the family. |
| 3 | Substituted | The government purchased the apartment block from the family. |
| 3 | Substituted | The company leased the apartment block to the family. |
| 3 | Substituted | The company purchased a valuable painting from the family. |
| 3 | Modified | In a calculated manoeuvre the company swiftly purchased the apartment block from the family. |
| 3 | Modified | A small company purchased the drab concrete apartment block from the family for later redevelopment. |
| 3 | Modified | Shortly before the election the company unwittingly purchased a defective apartment block from the family. |
| 3 | Modified | The foreign company purchased a decrepit apartment block from the wealthy family for an outrageous price. |
| 3 | Swapped | The family blocked the purchase of the apartment by the company. |
| 3 | Swapped | The family blocked the purchase of the apartment building by the company. |
| 3 | Swapped | The family blocked the purchase of their vintage car by the company. |
| 3 | Swapped | The family secured the purchase of their apartment by the company. |
| 3 | Swapped | The government blocked the purchase of the apartment by the company. |
| 3 | Swapped | The stubborn family blocked the purchase of the apartment by the company for six months. |
| 3 | Swapped | The family from across town blocked the purchase of the modern apartment by the unpopular company. |
| 3 | Swapped | After a long court battle the family finally blocked the purchase of the apartment by the unscrupulous corporation. |
| 3 | Swapped | For no good reason the family blocked the purchase of an old apartment by the local company for six months. |
| 4 | Base | The dog smelled the strange object near the children. |
| 4 | Same | The dog sniffed the strange object near the children. |
| 4 | Substituted | The cat smelled the strange object near the children. |
| 4 | Substituted | The dog noticed the strange object near the children. |

| | | |
|---|---|---|
| 4 | Substituted | The dog smelled smoke near the children. |
| 4 | Modified | The dog cautiously sniffed the mysterious object adjacent to the group of children. |
| 4 | Modified | The cute little dog smelled a strange object near the young children in the grass. |
| 4 | Modified | The feral dog smelled a strange rusty object near the children at the beach. |
| 4 | Modified | With its heightened sensory perception the dog smelled an unusual object near the large group of children at the school. |
| 4 | Swapped | The children objected to the unpleasant smell of the dog. |
| 4 | Swapped | The children objected to the dog's disagreeable smell. |
| 4 | Swapped | The father objected to the unpleasant smell of the dog. |
| 4 | Swapped | The children noticed the unpleasant smell of the dog. |
| 4 | Swapped | The children objected to the unpleasant smell of the cat. |
| 4 | Swapped | The children rarely objected to the unpleasant smell of their neighbour's dog. |
| 4 | Swapped | The group of children objected to the unpleasant smell of the large black dog from the park. |
| 4 | Swapped | The uncaring children objected to the smell of the sick dog at the vet. |
| 4 | Swapped | After five long days the disappointed children finally objected to the unpleasant smell of their favourite new dog. |
| 5 | Base | The lobbyist wrote to the politician about the reporter. |
| 5 | Same | The lobbyist wrote to the politician about the journalist. |
| 5 | Substituted | The nonprofit organisation wrote to the politician about the reporter. |
| 5 | Substituted | The lobbyist warned the politician about the reporter. |
| 5 | Substituted | The lobbyist wrote to the newspaper about the reporter. |
| 5 | Modified | In response to recent events the lobbyist quickly wrote to the politician about the reporter. |
| 5 | Modified | Over the weekend the lobbyist wrote a comprehensive report to the politician about the disgraced reporter. |
| 5 | Modified | The experienced lobbyist wrote a painstakingly detailed email to the politician about the reporter's recent activities. |
| 5 | Modified | The lobbyist composed an eloquent message to the politician on behalf of his client about the reporter. |
| 5 | Swapped | The reporter wrote to the lobbyist about the politician. |
| 5 | Swapped | The journalist wrote to the lobbyist about the politician. |
| 5 | Swapped | The academic wrote to the lobbyist about the politician. |
| 5 | Swapped | The reporter interviewed the lobbyist about the politician. |
| 5 | Swapped | The reporter wrote to the nonprofit organisation about the politician. |
| 5 | Swapped | On her new laptop the reporter wrote a short blog post for the lobbyist regarding the politician. |
| 5 | Swapped | The reporter regularly wrote to the lobbyist about the powerful politician via an encrypted message system. |
| 5 | Swapped | Using pen and paper the reporter meticulously wrote a personal letter to the lobbyist about the politician. |
| 5 | Swapped | Out of concern for the public, the reporter wrote to the lobbyist about the politician's recent decisions. |
| 6 | Base | The shopkeeper displayed the watch for customers. |
| 6 | Same | The shopkeeper displayed the watch for his customers. |
| 6 | Substituted | The salesperson displayed the watch for customers. |
| 6 | Substituted | The shopkeeper marketed the watch to customers. |
| 6 | Substituted | The shopkeeper displayed books for customers. |
| 6 | Modified | Each day the shopkeeper displayed the watch to customers in his store. |
| 6 | Modified | During busy periods the shopkeeper prominently displayed the watch for customers. |
| 6 | Modified | The incompetent shopkeeper foolishly displayed the watch to customers in an obscure location. |

| | | |
|---|---|---|
| 6 | Modified | As a marketing technique the shopkeeper prominently placed the watch on a dedicated stand for customers. |
| 6 | Swapped | The customer watched over the display for the shopkeeper. |
| 6 | Swapped | A customer watched over the display for the shopkeeper. |
| 6 | Swapped | The surveillance camera watched over the display for the shopkeeper. |
| 6 | Swapped | The customer critiqued the display for the shopkeeper. |
| 6 | Swapped | The customer watched over the model for the shopkeeper. |
| 6 | Swapped | Every day the customer watched over the display for the shopkeeper during lunch. |
| 6 | Swapped | The customer patiently watched over the well-organized display for the shopkeeper for hours on end. |
| 6 | Swapped | The unscrupulous customer greedily watched the display behind the shopkeeper's back during the busiest time of day. |
| 6 | Swapped | In the back room the nosey customer watched over the goods display for the shopkeeper on the security monitor. |

#### 1.3 Supplementary figures

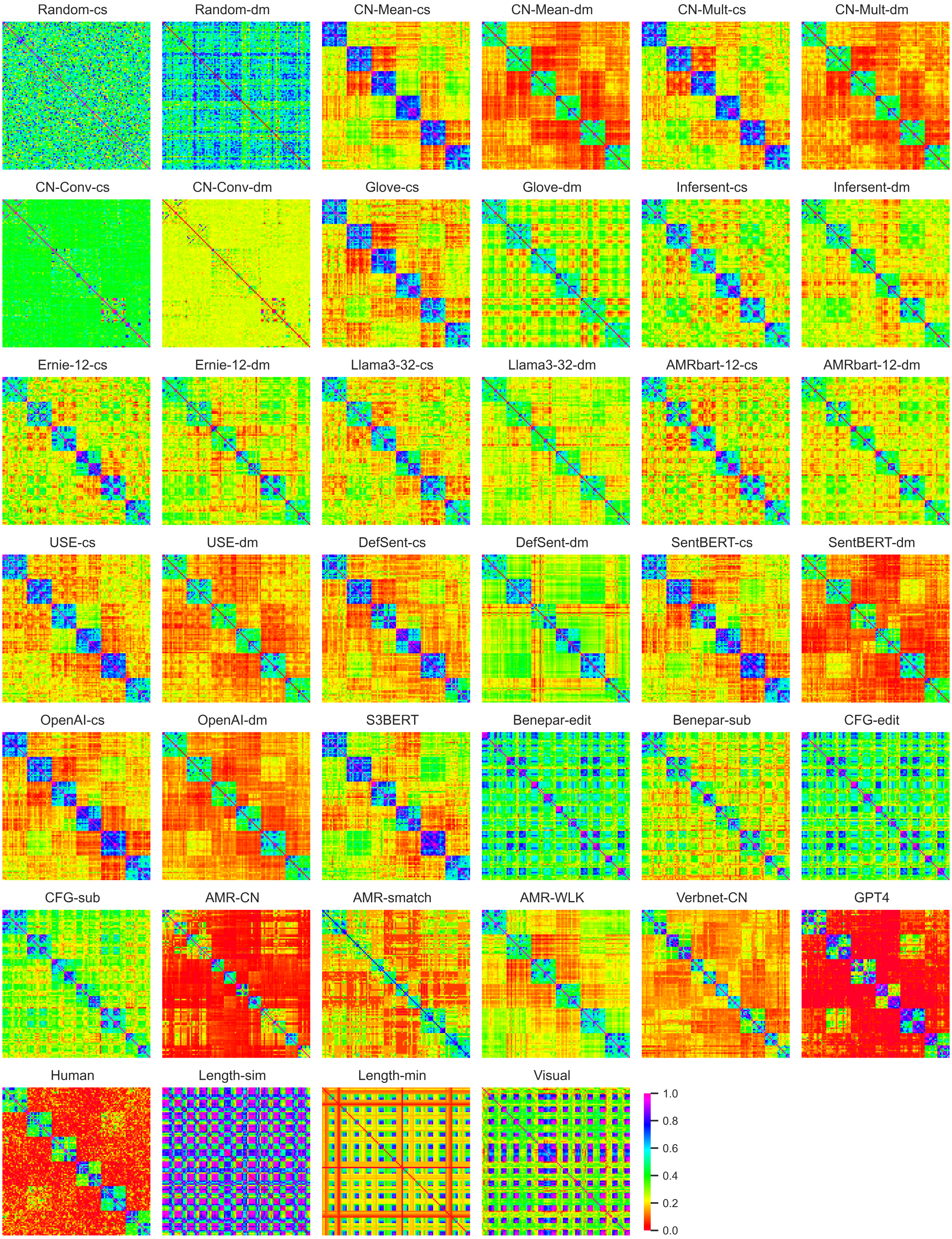


**Fig. S1: RSA matrices for all computational models.** Representational similarity matrices for all 21 models, as well as human and GPT-4 ratings, and three control variables related to sentence lenth and visual similarity. For vector-based models, two similarity metrics were used: cosine similarity ('cs') and DIEM similarity ('dm'). Block diagonal patterns are visible for all models aside from random embeddings, indicating all models are sensitive to lexical similarity. Most transformers show only limited similarity to sentence structure, as evident from the lack of distinction between off-diagonal 'swapped' sentence pairs and on-diagonal 'modified' and 'substituted' sentence pairs. By contrast, the Graph-based and the VerbNet-CN hybrid models generally show greater sensitivity to such structural differences.

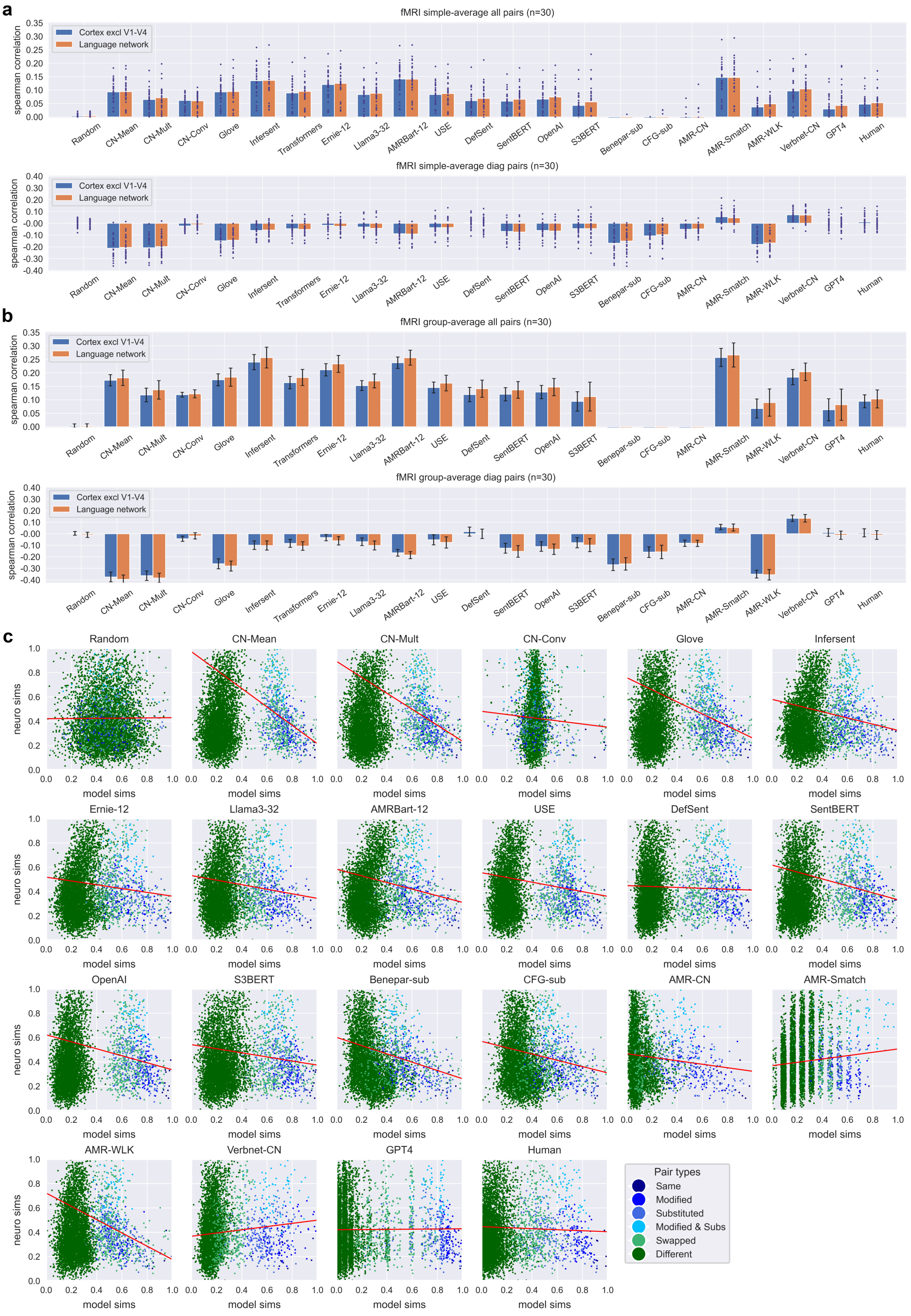


**Fig. S2: Correlations between all computational models and brain activity.** Partial correlations between RSA matrices of all computational models and the brain RSA matrix, controlling for differences in sentence length. Blue bars show correlations computed over all stable (excluding visual regions V1-V4), while green bars show correlations for stable voxels in the language network. **a)** Partial correlations for each individual participant shown as blue dots, with the simple average over individual correlations shown as a bar. **b)** Partial correlations computed using the group-averaged RSA matrix. Error bars show 95% confidence intervals calculated by bootstrap resampling over participants. **c)** Scatterplots showing the relationship between model similarities (horizontal axis) and group-average neural similarities (vertical axis) for all computational models. Each dot corresponds to a single pairwise similarity, scaled to between 0 and 1 for visualisation. While all sentence pairs are shown, regression lines (red) are computed over the block diagonal pairs only.

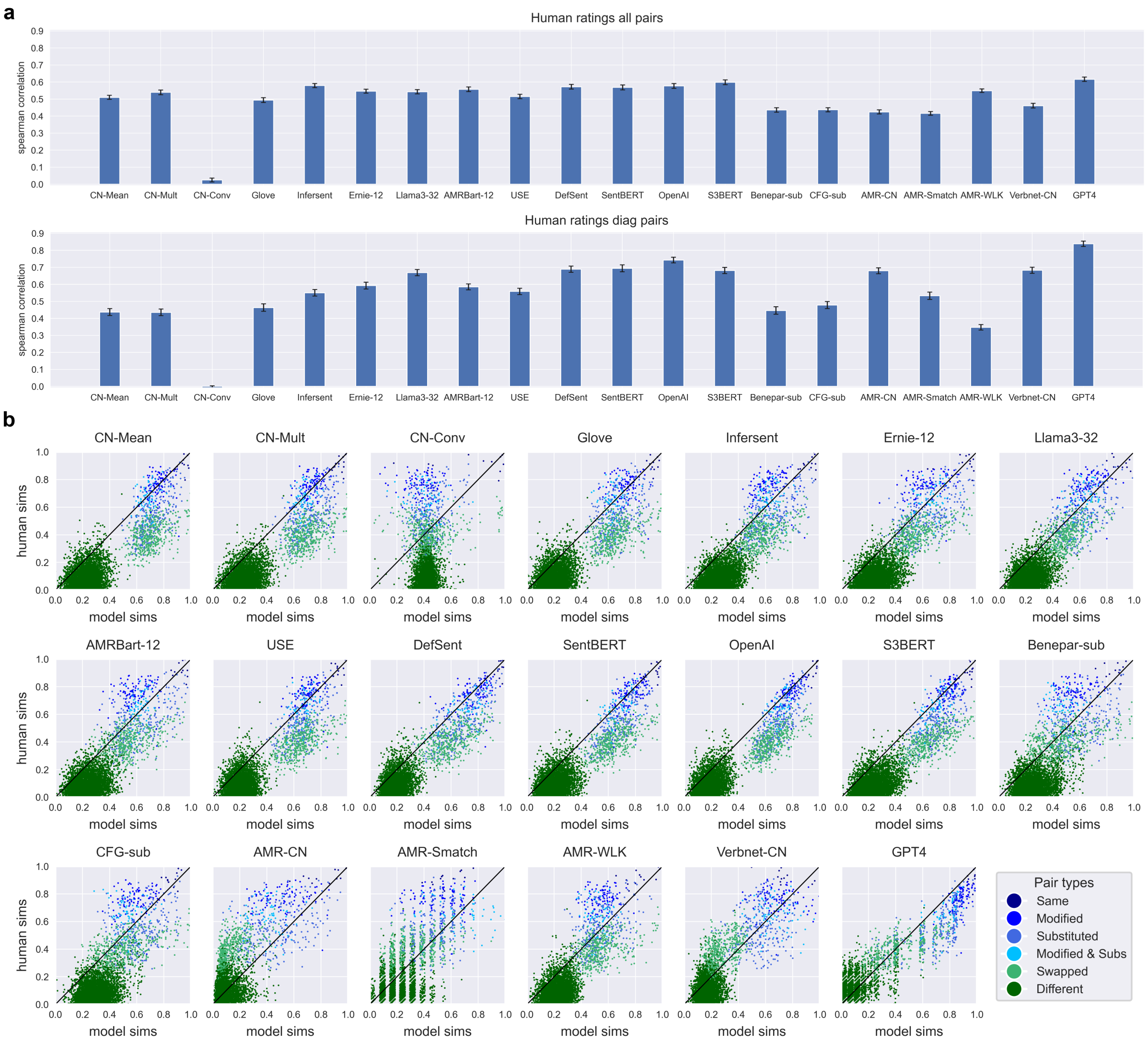


**Fig. S3: Correlations between behavioural ratings of sentence similarity and model similarities for all computational models.** **a)** Correlations between RSA matrices of all computational models and human-rated similarities computed over all sentence pairs (top) and block diagonal sentence pairs (bottom). Error bars show 95% confidence intervals calculated by bootstrap resampling over raters. **b)** Scatterplots showing the relationship between model similarities (horizontal axis) and human rated similarities (vertical axis) for all computational models. Each dot corresponds to a single pairwise similarity. The 45-degree line (black) shows a hypothetical line of perfect fit between model and human similarities.

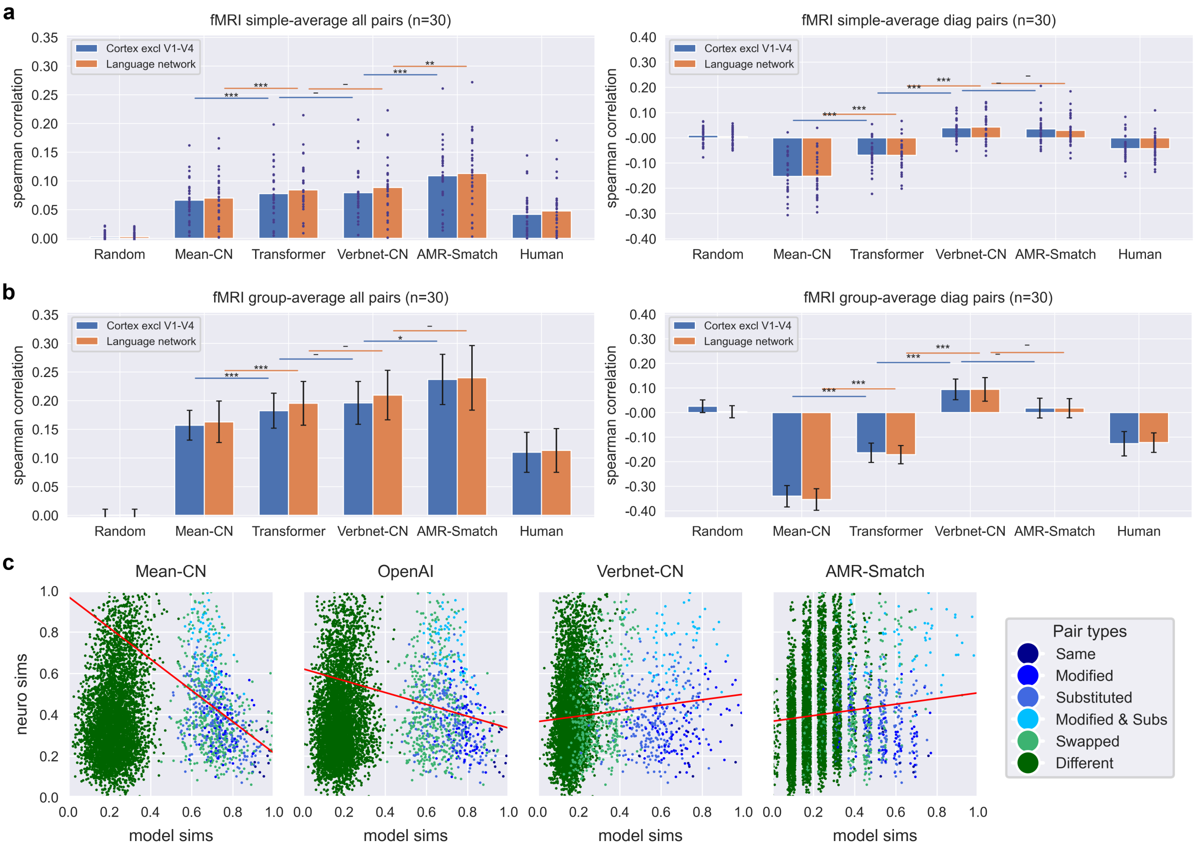


**Fig. S4: Model correlations with brain activity for all sentence pairs and the block-diagonal subset of sentence pairs, with an additional control for minimum sentence length.** Partial correlations between RSA matrices of five computational models and the brain RSA matrix, controlling for differences in sentence length and minimum sentence length. 'Human' refers to behavioural ratings. Blue bars show correlations computed over all stable (excluding visual regions V1-V4), while green bars show correlations for stable voxels in the language network. Notation for statistical significance: * for p<0.05, ** for p<0.01, and *** for p<0.001, with Bonferroni correction for three independent comparisons. **a)** Partial correlations for each individual participant shown as blue dots, with the simple average over individual correlations shown as a bar. **b)** Partial correlations computed using the group-averaged RSA matrix. Error bars show 95% confidence intervals calculated by bootstrap resampling over participants. **c)** Scatterplots showing the relationship between model similarities (horizontal axis) and group-average neural similarities (vertical axis) for all four computational models. Each dot corresponds to a single pairwise similarity after regressing out the effects of sentence length similarity and minimum sentence length. Note that the displayed correlation for the AMR-Smatch is inaccurate owing the discrete nature of SMATCH similarities, which leads to inconsistencies when using regression residuals to estimate brain similarities after controlling for length differences and the minimum sentence length. Similarities have been rescaled to between 0 and 1 for visualisation. While all sentence pairs are shown, regression lines (red) are computed over the block diagonal pairs only.

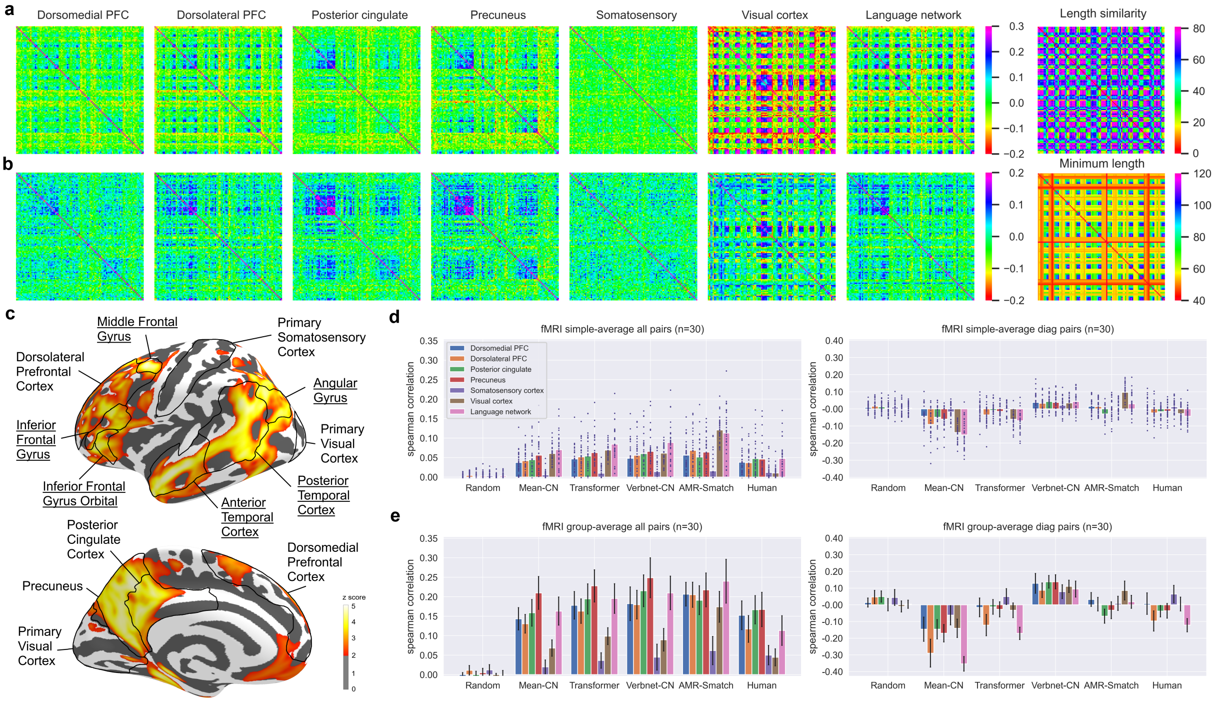


**Fig. S5: Comparison of sentence representations and model correlations across brain regions, with an additional control for minimum sentence length.** **a)** Representational similarity matrices for various cortical regions, computed controlling for differences in sentence length and minimum sentence length. **b)** Representational similarity matrices for various cortical regions, computed controlling for difference in sentence length and minimum sentence length. Block diagonal structure is visible for all regions outside the visual cortex. **c)** Searchlight RSA for Hybrid model using 8mm radius showing cortical regions of interest, with those part of the language network underlined. RSA correlations are thresholded at z=2. **d)** Partial correlations controlling for differences in sentence length and minimum sentence length by cortical region, with the simple average over individual correlations shown as a bar. Error bars show 95% confidence intervals calculated by bootstrap resampling over participants. **e)** Partial correlations controlling for differences in sentence length and minimum sentence length computed using the group-averaged RSA matrix, shown by cortical region. Error bars show 95% confidence intervals calculated by bootstrap resampling over participants.

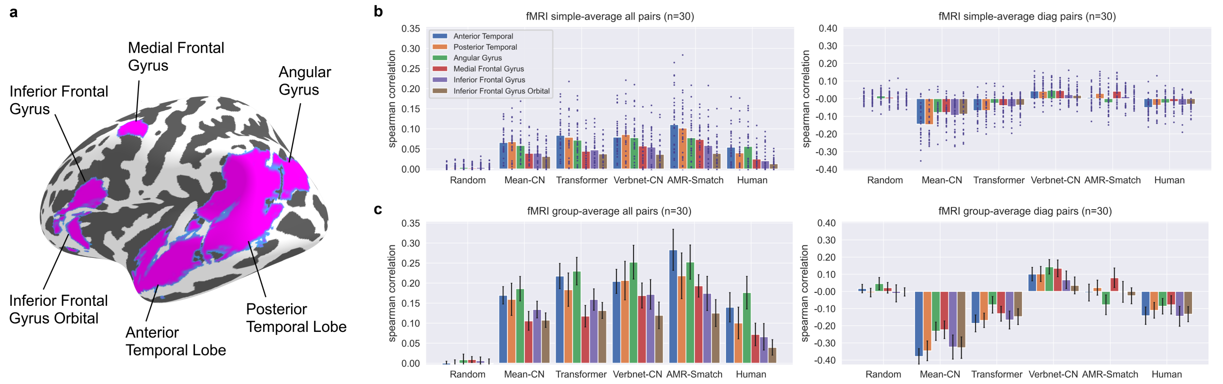


**Fig. S6: Comparison of model correlations across subregions of the language network, with an additional control for minimum sentence length.** **a)** Regions within the language network. **b)** Partial correlations controlling for differences in sentence length and minimum sentence length shown by language network subregion, with each individual participant shown as blue dots, and the simple average over individual correlations shown as a bar. **c)** Partial correlations controlling for differences in sentence length and minimum sentence length computed using the group-averaged RSA matrix, shown by language network subregion. Error bars show 95% confidence intervals calculated by bootstrap resampling over participants.

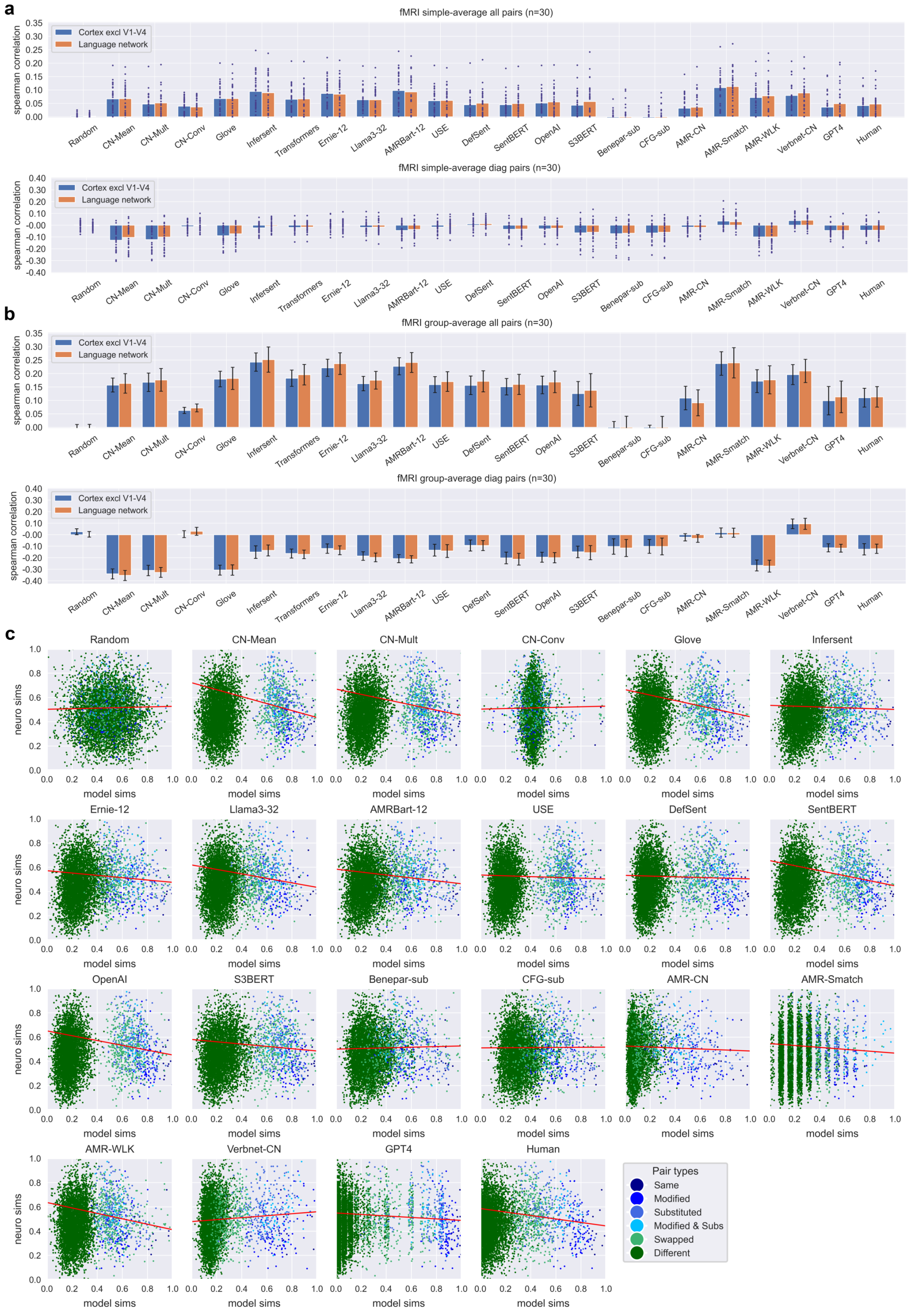


**Fig. S7: Correlations between all computational models and brain activity, with an additional control for minimum sentence length.** Partial correlations between RSA matrices of all computational models and the brain RSA matrix, controlling for differences in sentence length and minimum sentence length. Blue bars show correlations computed over all stable (excluding visual regions V1-V4), while green bars show correlations for stable voxels in the language network. **a)** Partial correlations for each individual participant shown as blue dots, with the simple average over individual correlations shown as a bar. **b)** Partial correlations computed using the group-averaged RSA matrix. Error bars show 95% confidence intervals calculated by bootstrap resampling over participants. **c)** Scatterplots showing the relationship between model similarities (horizontal axis) and group-average neural similarities (vertical axis) for all computational models. Each dot corresponds to a single pairwise similarity, scaled to between 0 and 1 for visualisation. While all sentence pairs are shown, regression lines (red) are computed over the block diagonal pairs only.

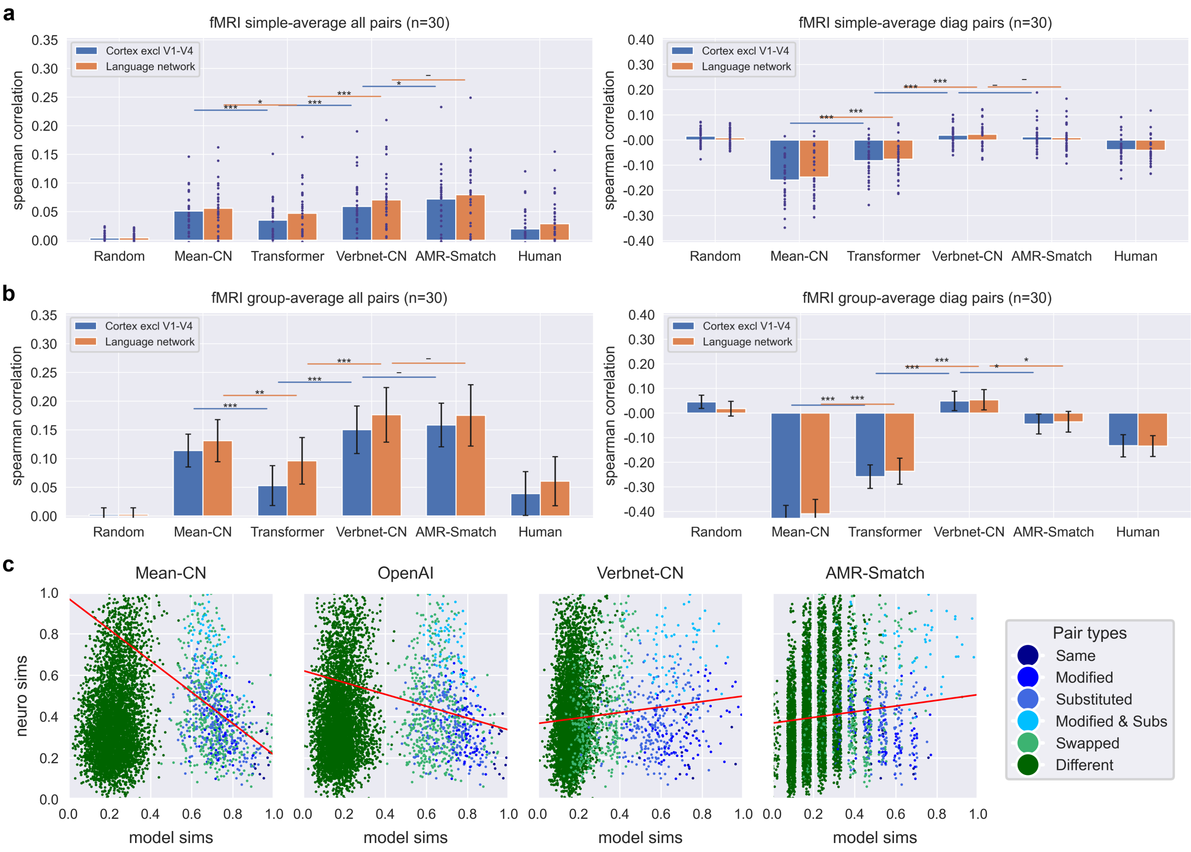


**Fig. S8: Model correlations with brain activity for all sentence pairs and the block-diagonal subset of sentence pairs, with visual similarity control.** Partial correlations between RSA matrices of five computational models and the brain RSA matrix, controlling for the RSA similarity of visual cortex activity averaged over participants. 'Human' refers to behavioural ratings. Blue bars show correlations computed over all stable (excluding visual regions V1-V4), while green bars show correlations for stable voxels in the language network. Notation for statistical significance: * for $p<0.05$, ** for $p<0.01$, and *** for $p<0.001$, with Bonferroni correction for three independent comparisons. **a)** Partial correlations for each individual participant shown as blue dots, with the simple average over individual correlations shown as a bar. **b)** Partial correlations computed using the group-averaged RSA matrix. Error bars show 95% confidence intervals calculated by bootstrap resampling over participants. **c)** Scatterplots showing the relationship between model similarities (horizontal axis) and group-average neural similarities (vertical axis) for all four computational models. Each dot corresponds to a single pairwise similarity after regressing out the subject-averaged visual cortex similarities. Note that the displayed correlation for the AMR-Smatch is inaccurate owing the discrete nature of SMATCH similarities, which leads to inconsistencies when using regression residuals to estimate brain similarities. Similarities have been rescaled to between 0 and 1 for visualisation. While all sentence pairs are shown, regression lines (red) are computed over the block diagonal pairs only.

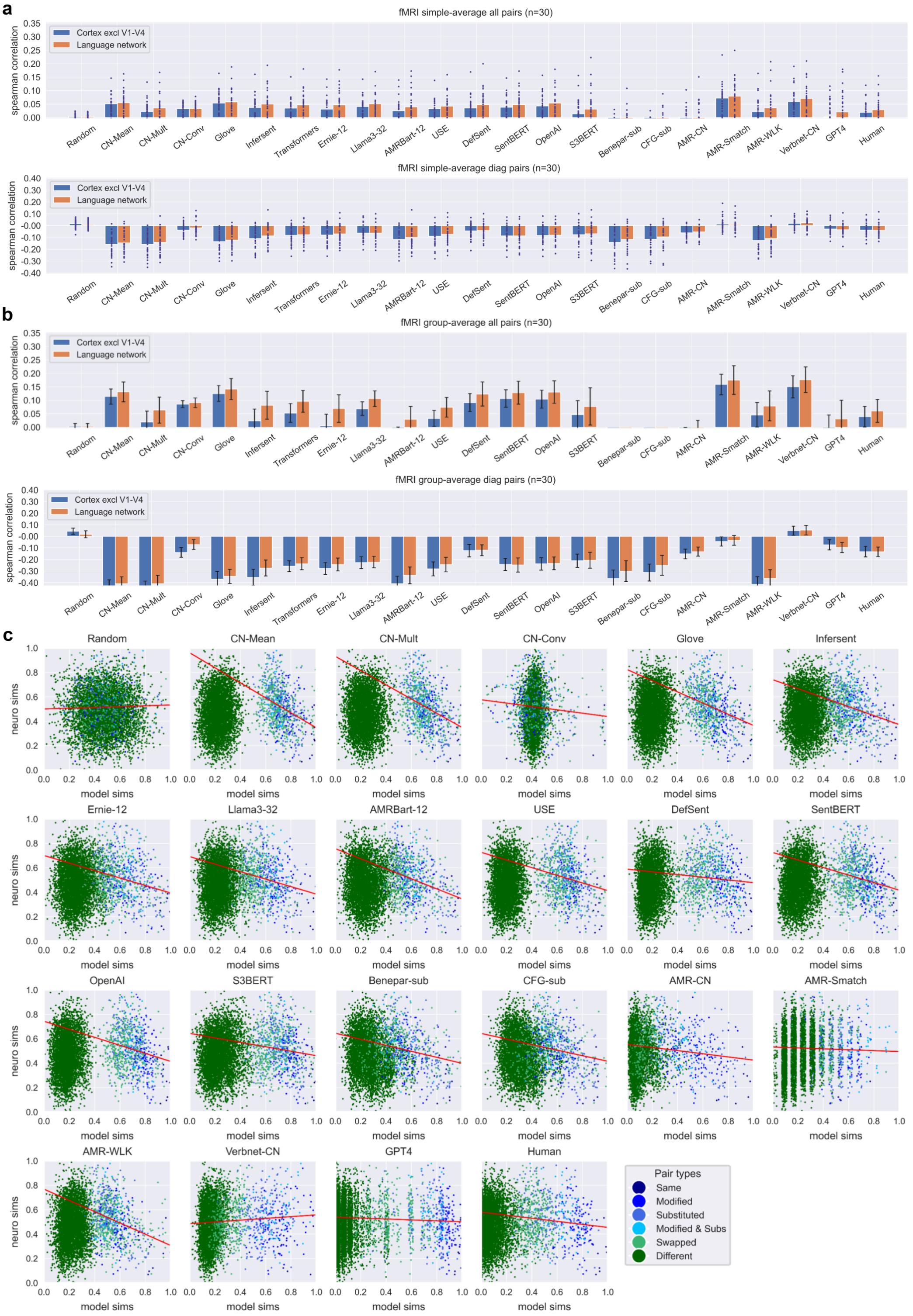


**Fig. S9: Correlations between all computational models and brain activity, with visual similarity control.** Partial correlations between RSA matrices of all computational models and the brain RSA matrix, controlling for the RSA similarity of visual cortex activity averaged over participants. Blue bars show correlations computed over all stable (excluding visual regions V1-V4), while green bars show correlations for stable voxels in the language network. **a)** Partial correlations for each individual participant shown as blue dots, with the simple average over individual correlations shown as a bar. **b)** Partial correlations computed using the group-averaged RSA matrix. Error bars show 95% confidence intervals calculated by bootstrap resampling over participants. **c)** Scatterplots showing the relationship between model similarities (horizontal axis) and group-average neural similarities (vertical axis) for all computational models. Each dot corresponds to a single pairwise similarity, scaled to between 0 and 1 for visualisation. While all sentence pairs are shown, regression lines (red) are computed over the block diagonal pairs only.

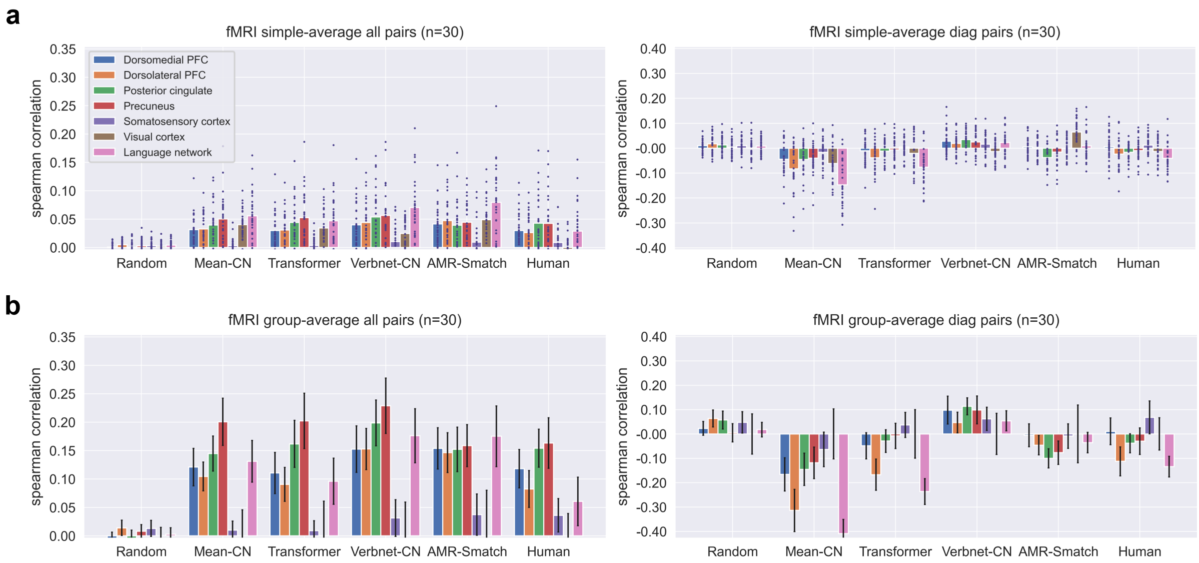


**Fig. S10: Comparison of sentence representations and model correlations across brain regions, with visual similarity control**. **a)** Partial correlations controlling for differences in sentence length and minimum sentence length by cortical region, with the simple average over individual correlations shown as a bar. Error bars show 95% confidence intervals calculated by bootstrap resampling over participants. **b)** Partial correlations controlling for differences in sentence length and minimum sentence length computed using the group-averaged RSA matrix, shown by cortical region. Error bars show 95% confidence intervals calculated by bootstrap resampling over participants.

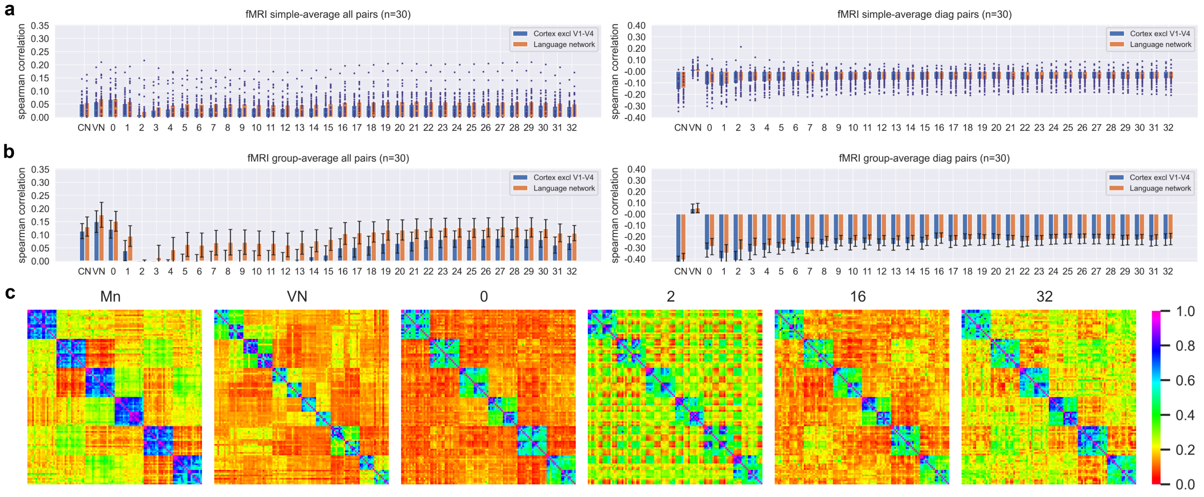


**Fig. S11: Correlations between individual fMRI data and Llama 3 layers, with visual similarity control.** Average correlations between RSA matrices of each layer of Llama 3 and brain RSA matrix of each participant, controlling for the RSA similarity of visual cortex activity averaged over participants. Mean-CN (CN) and VerbNet-CN hybrid (VN) models are also shown for comparison. **a)** Partial correlations for each individual participant shown as blue dots, with the simple average over individual correlations shown as a bar. **b)** Partial correlations computed using the group-averaged RSA matrix. Error bars show 95% confidence intervals calculated by bootstrap resampling over participants. **c)** RSA matrices for the Mean-CN and VerbNet-CN models, along with selected layers of Llama 3, computed controlling for visual similarity.

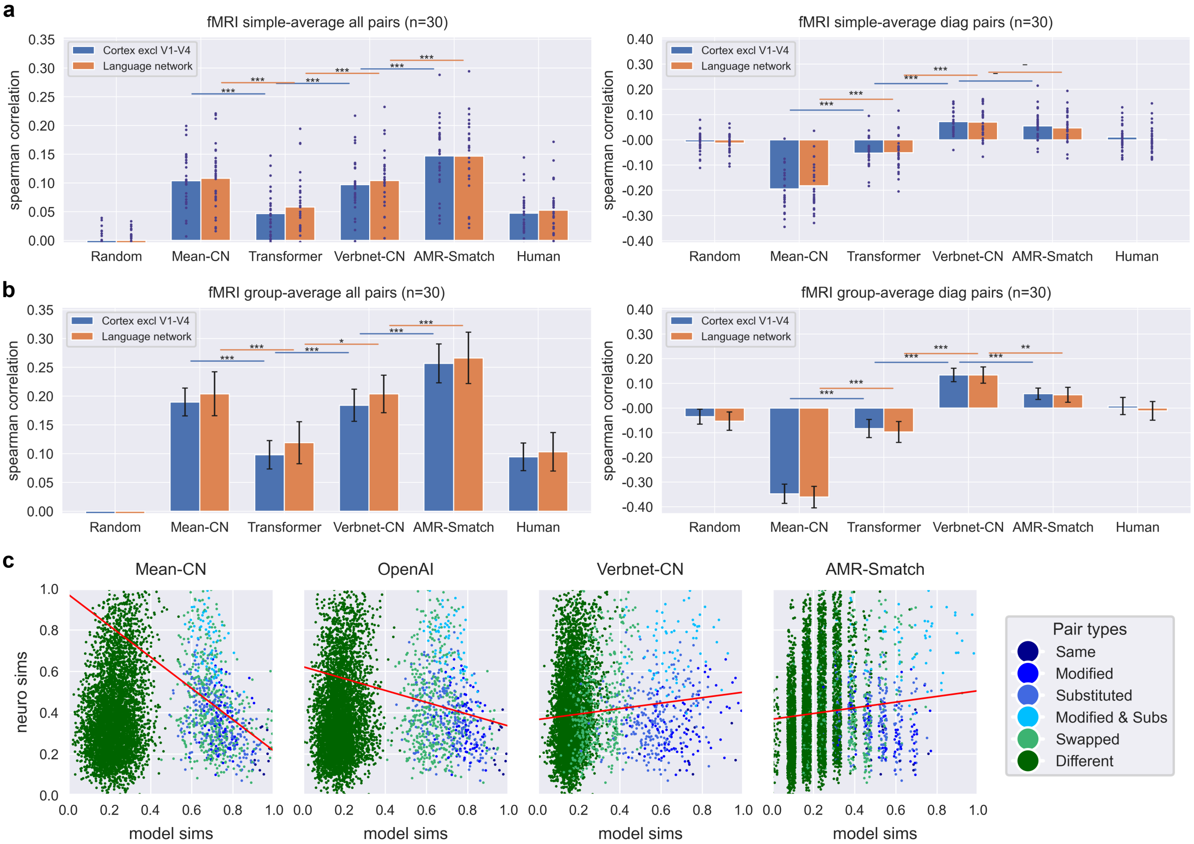


**Fig. S12: Model correlations with brain activity for all sentence pairs and the block-diagonal subset of sentence pairs using the DIEM similarity metric.** Partial correlations between RSA matrices of five computational models and the brain RSA matrix, controlling for differences in sentence length. 'Human' refers to behavioural ratings. Blue bars show correlations computed over all stable (excluding visual regions V1-V4), while green bars show correlations for stable voxels in the language network. Notation for statistical significance: * for p<0.05, ** for p<0.01, and *** for p<0.001, with Bonferroni correction for three independent comparisons. **a)** Partial correlations for each individual participant shown as blue dots, with the simple average over individual correlations shown as a bar. **b)** Partial correlations computed using the group-averaged RSA matrix. Error bars show 95% confidence intervals calculated by bootstrap resampling over participants. **c)** Scatterplots showing the relationship between model similarities (horizontal axis) and group-average neural similarities (vertical axis) for all four computational models. Each dot corresponds to a single pairwise similarity after regressing out the effects of sentence length similarity and minimum sentence length. Note that the displayed correlation for the AMR-Smatch is inaccurate owing the discrete nature of SMATCH similarities, which leads to inconsistencies when using regression residuals to estimate brain similarities after controlling for length differences and the minimum sentence length. Similarities have been rescaled to between 0 and 1 for visualisation. While all sentence pairs are shown, regression lines (red) are computed over the block diagonal pairs only.

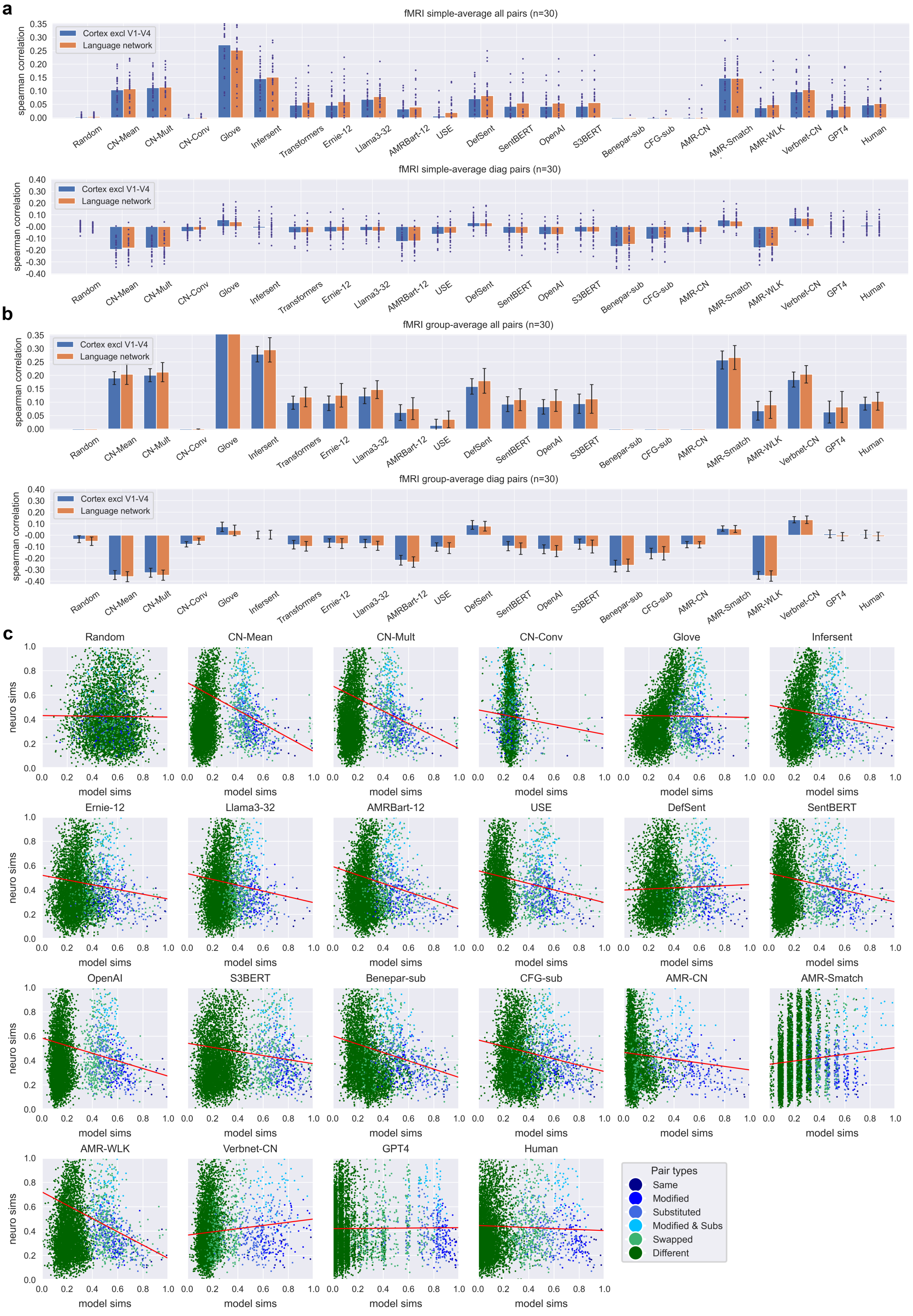


**Fig. S13: Correlations between all computational models and brain activity using DIEM similarity metric.** Partial correlations between RSA matrices of all computational models and the brain RSA matrix, controlling for differences in sentence length. Blue bars show correlations computed over all stable (excluding visual regions V1-V4), while green bars show correlations for stable voxels in the language network. **a)** Partial correlations for each individual participant shown as blue dots, with the simple average over individual correlations shown as a bar. **b)** Partial correlations computed using the group-averaged RSA matrix. Error bars show 95% confidence intervals calculated by bootstrap resampling over participants. **c)** Scatterplots showing the relationship between model similarities (horizontal axis) and group-average neural similarities (vertical axis) for all computational models. Each dot corresponds to a single pairwise similarity, scaled to between 0 and 1 for visualisation. While all sentence pairs are shown, regression lines (red) are computed over the block diagonal pairs only.

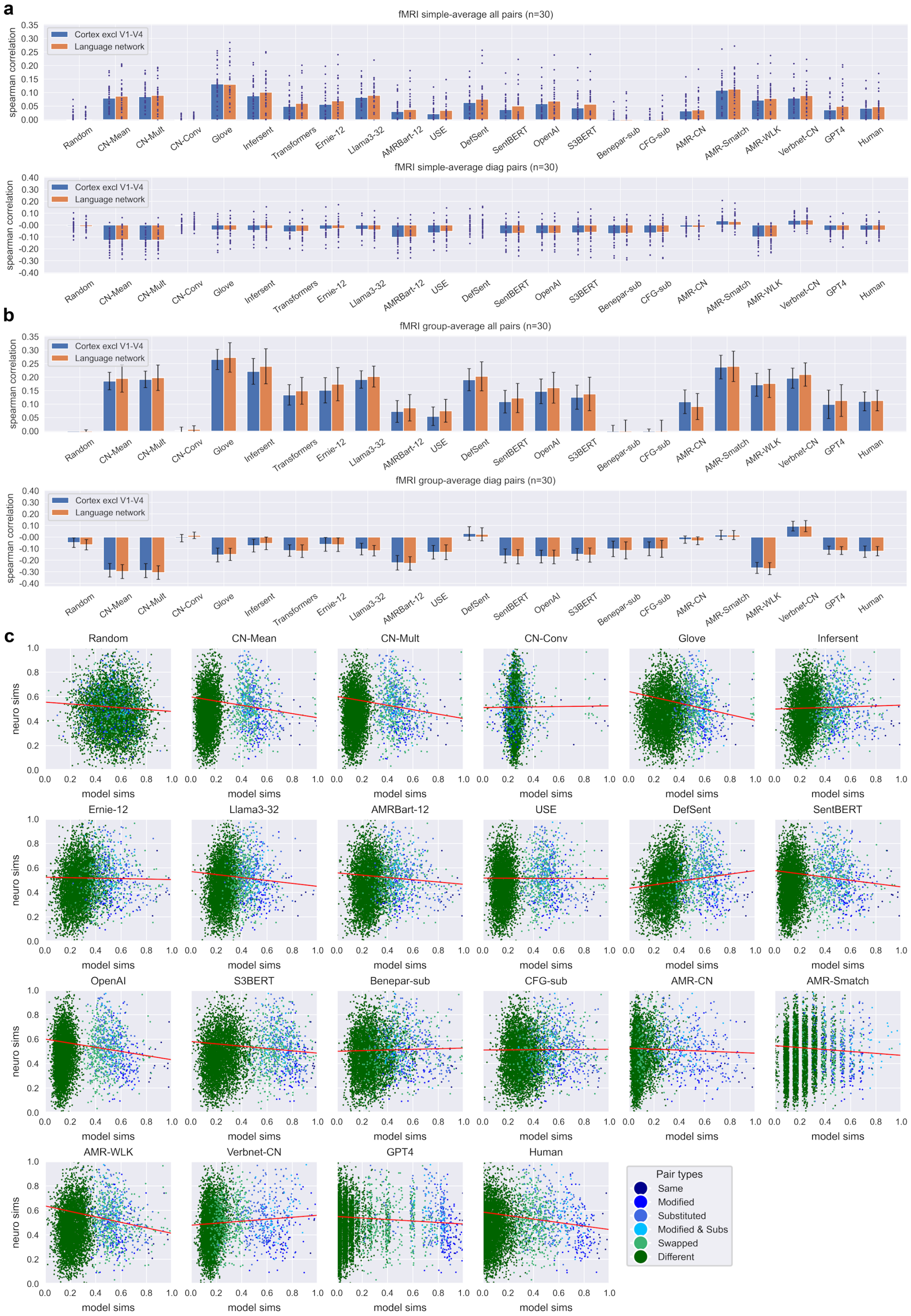


**Fig. S14: Correlations between all computational models and brain activity using DIEM similarity metric, with an additional control for minimum sentence length.** Partial correlations between RSA matrices of all computational models and the brain RSA matrix, controlling for differences in sentence length and minimum sentence length. Blue bars show correlations computed over all stable (excluding visual regions V1-V4), while green bars show correlations for stable voxels in the language network. **a)** Partial correlations for each individual participant shown as blue dots, with the simple average over individual correlations shown as a bar. **b)** Partial correlations computed using the group-averaged RSA matrix. Error bars show 95% confidence intervals calculated by bootstrap resampling over participants. **c)** Scatterplots showing the relationship between model similarities (horizontal axis) and group-average neural similarities (vertical axis) for all computational models. Each dot corresponds to a single pairwise similarity, scaled to between 0 and 1 for visualisation. While all sentence pairs are shown, regression lines (red) are computed over the block diagonal pairs only.

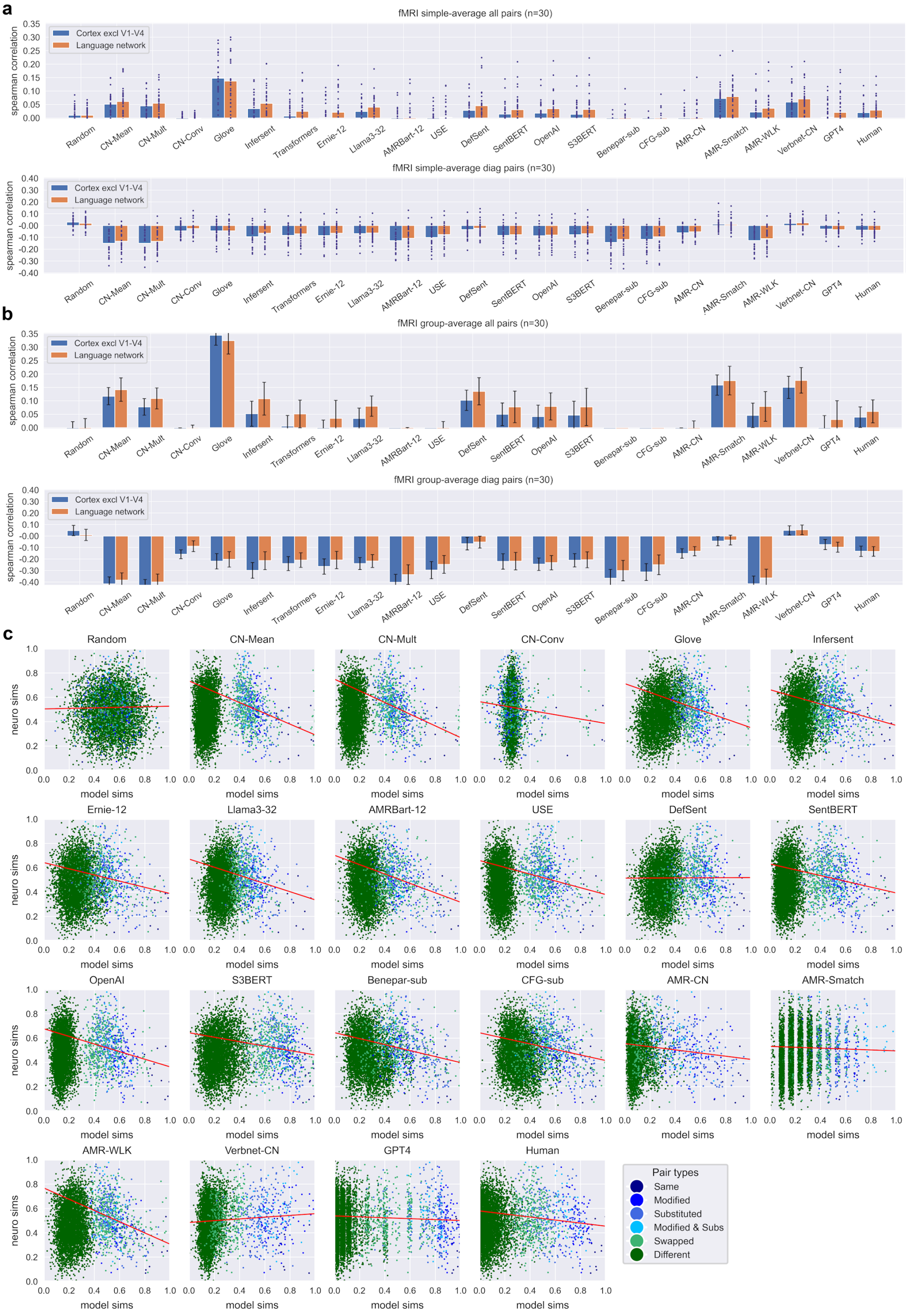


**Fig. S15: Correlations between all computational models and brain activity using DIEM similarity metric, with visual similarity control.** Partial correlations between RSA matrices of all computational models and the brain RSA matrix, controlling for the RSA similarity of visual cortex activity averaged over participants. Blue bars show correlations computed over all stable (excluding visual regions V1-V4), while green bars show correlations for stable voxels in the language network. **a)** Partial correlations for each individual participant shown as blue dots, with the simple average over individual correlations shown as a bar. **b)** Partial correlations computed using the group-averaged RSA matrix. Error bars show 95% confidence intervals calculated by bootstrap resampling over participants. **c)** Scatterplots showing the relationship between model similarities (horizontal axis) and group-average neural similarities (vertical axis) for all computational models. Each dot corresponds to a single pairwise similarity, scaled to between 0 and 1 for visualisation. While all sentence pairs are shown, regression lines (red) are computed over the block diagonal pairs only.

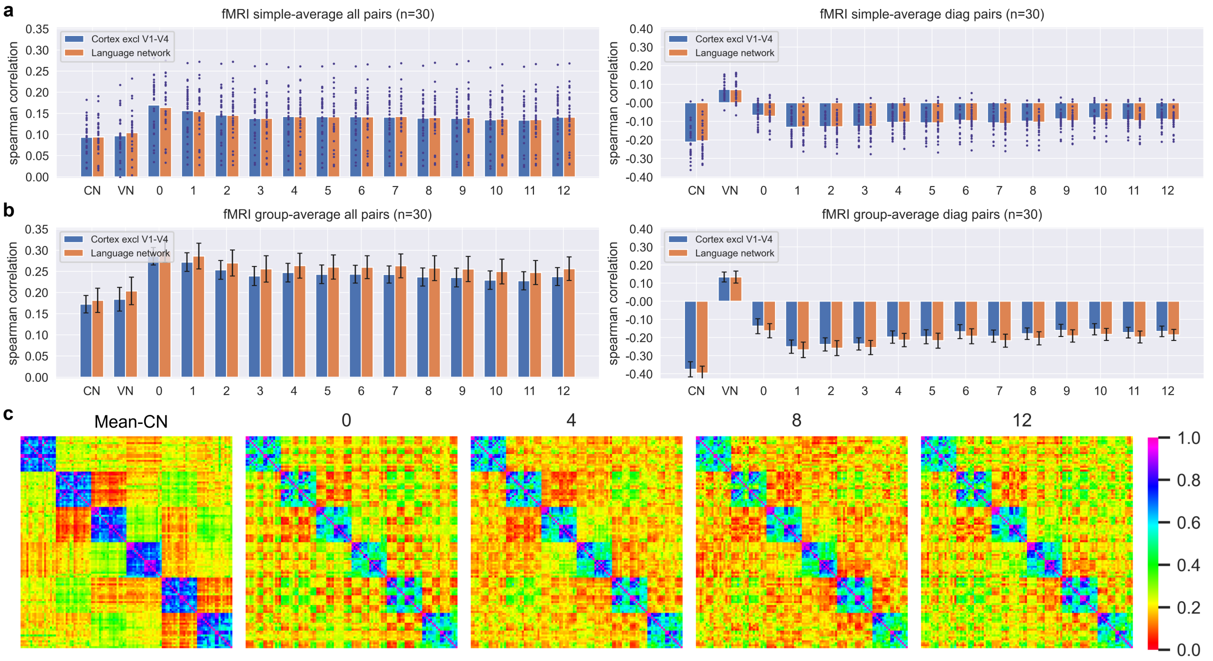


**Fig. S16: Average correlations between RSA matrices of each layer of AMRBart and the brain RSA matrix of each participant**. Mean-CN (Mn) and VerbNet-CN hybrid (VN) models are also shown for comparison. **a)** Partial correlations for each individual participant shown as blue dots, with the simple average over individual correlations shown as a bar. **b)** Partial correlations computed using the group-averaged RSA matrix. Error bars show 95% confidence intervals calculated by bootstrap resampling over participants. **c)** RSA matrices for the Mean-CN and VerbNet-CN models, along with selected layers of the AMRBart.

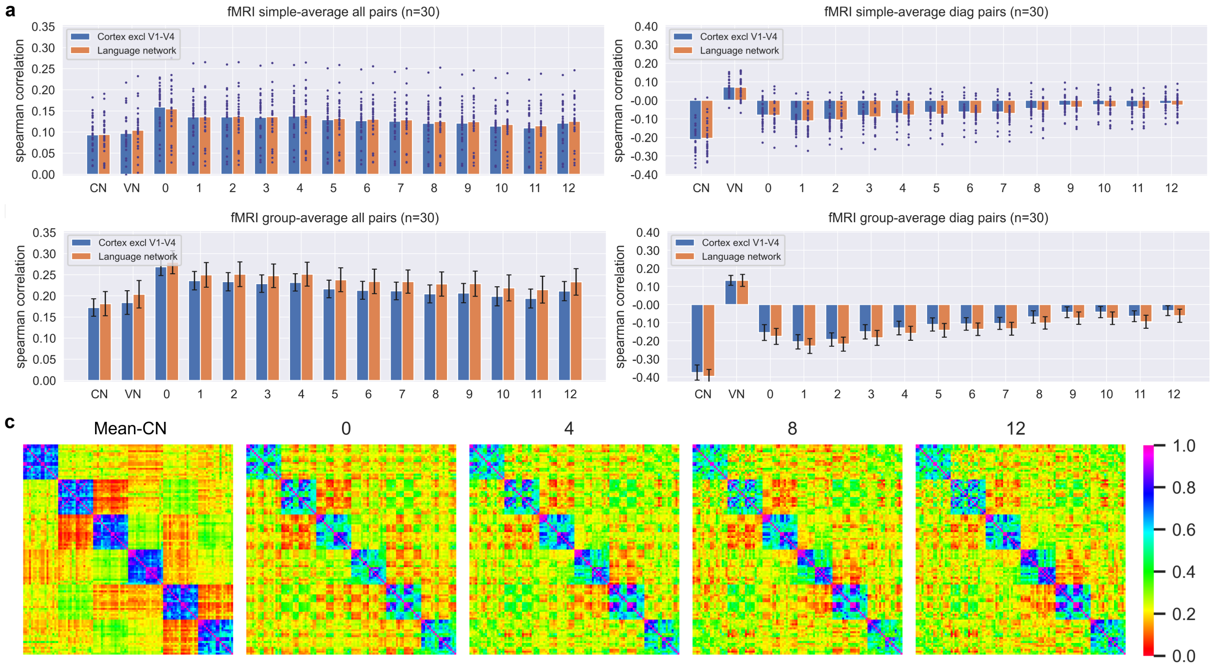


**Fig. S17: Average correlations between RSA matrices of each layer of ERNIE 2.0 and brain RSA matrix of each participant**. Mean-CN (Mn) and VerbNet-CN hybrid (VN) models are also shown for comparison. **a)** Partial correlations for each individual participant shown as blue dots, with the simple average over individual correlations shown as a bar. **b)** Partial correlations computed using the group-averaged RSA matrix. Error bars show 95% confidence intervals calculated by bootstrap resampling over participants. **c)** RSA matrices for the Mean-CN and VerbNet-CN models, along with selected layers of the ERNIE 2.0.

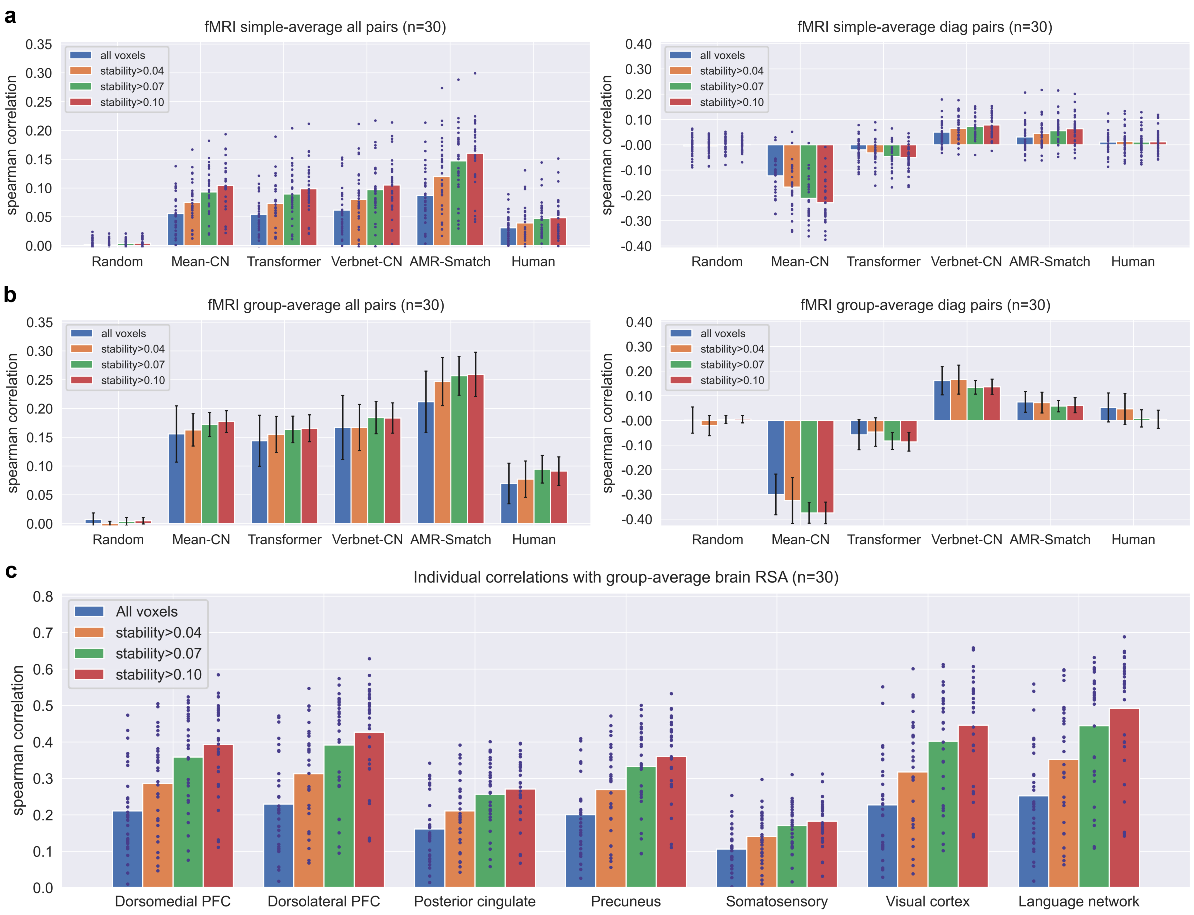


**Fig. S18: Comparison of model correlations with brain activity when varying the stability score threshold.** Partial correlations between RSA matrices of five computational models and the brain RSA matrix, controlling for differences in sentence length. Bars are colour-coded based on the threshold used for voxel inclusion. **a)** Partial correlations for each individual participant shown as blue dots, with the simple average over individual correlations shown as a bar. **b)** Partial correlations computed using the group-averaged RSA matrix. Error bars show 95% confidence intervals calculated by bootstrap resampling over participants. **c)** Correlations between RSA matrix of each individual participant and the group-averaged RSA matrix, organised by the voxel selection method used for computing the RSA matrix. Each dot corresponds to the correlation for a single participant.

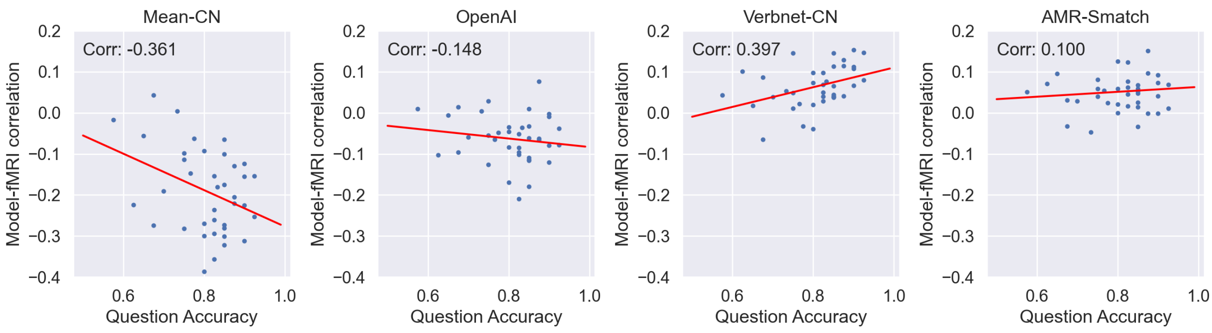


**Fig. S19: Correlation between participant accuracy in attention-check questions and model similarity to brain representations.** Scatterplot showing the accuracy score for each participant on attention check questions presented during the scanning run (horizontal axis) against the model correlation with all sentence pairs computed over all stable cortical voxels excluding V1-V4 (vertical axis). Each dot shows the results for a single participant, with all 38 participants included who achieved a score on the questions above chance level (25%).

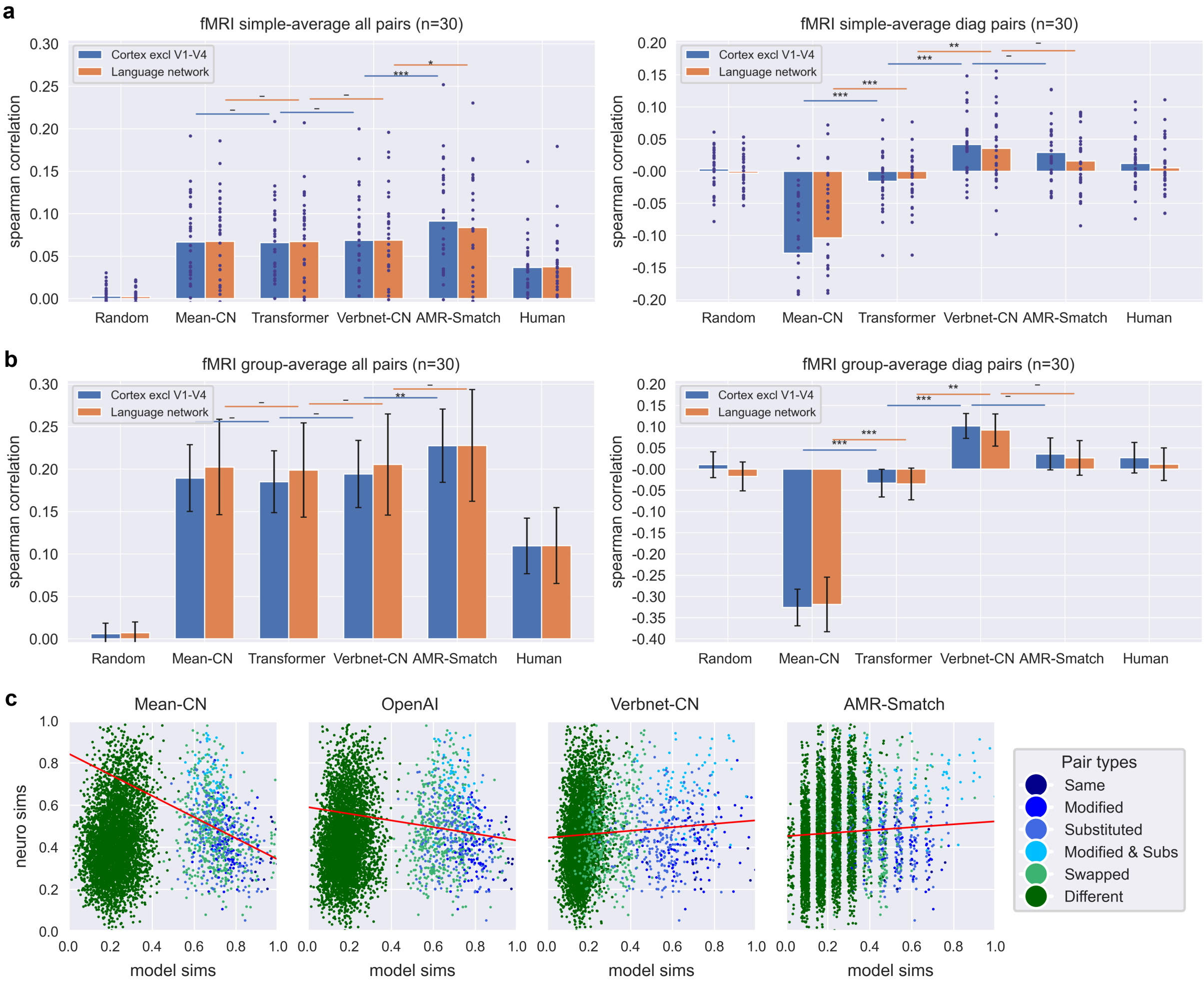


**Fig. S20: Model correlations with brain activity for all sentence pairs and the block-diagonal subset of sentence pairs, using the final 3s of each stimulus.** Partial correlations between RSA matrices of five computational models (Random, Mean, Transformers, Hybrid, and Graph) and the brain RSA matrix, controlling for differences in sentence length and using the final 3s for each stimulus. Blue bars show correlations computed over all stable (excluding visual regions V1-V4), while green bars show correlations for stable voxels in the language network. Notation for statistical significance: * for p<0.05, ** for p<0.01, and *** for p<0.001, with Bonferroni correction for three independent comparisons. **a)** Partial correlations for each individual participant shown as blue dots, with the simple average over individual correlations shown as a bar. **b)** Partial correlations computed using the group-averaged RSA matrix. Error bars show 95% confidence intervals calculated by bootstrap resampling over participants. **c)** Scatterplots showing the relationship between model similarities (horizontal axis) and group-average neural similarities (vertical axis) for all four computational models. Each dot corresponds to a single pairwise similarity, scaled to between 0 and 1 for visualisation. While all sentence pairs are shown, regression lines (red) are computed over the block diagonal pairs only.
